# Probing mechanical interaction of immune receptors and cytoskeleton by membrane nanotube extraction

**DOI:** 10.1101/2022.09.15.508080

**Authors:** Fabio Manca, Gautier Eich, Omar N’Dao, Lucie Normand, Kheya Sengupta, Laurent Limozin, Pierre-Henri Puech

## Abstract

The role of force application in immune cell recognition is now well established, the force being transmitted between the actin cytoskeleton to the anchoring ligands through receptors such as integrins. In this chain, the mechanics of the cytoskeleton to receptor link, though clearly crucial, remains poorly understood. To probe this link, we combine mechanical extraction of membrane tubes from T cells using optical tweezers, and fitting of the resulting force curves with a viscoelastic model taking into account the cell and relevant molecules. We solicit this link using four different antibodies against various membrane bound receptors: antiCD3 to target the T Cell Receptor (TCR) complex, antiCD45 for the long sugar CD45, and two clones of antiCD11 targeting open or closed conformation of LFA1 integrins. Upon disruption of the cytoskeleton, the stiffness of the link changes for two of the receptors, exposing the existence of a receptor to cytoskeleton link - namely TCR-complex and open LFA1, and does not change for the other two where no such a link was expected. Our integrated approach allows us to probe, for the first time, the mechanics of the intracellular receptor-cytoskeleton link in immune cells.

## Introduction

The importance of mechanics and mechanotransduction, at both molecular and cellular scales, is now well recognized in cell biology in general [Vogel and Sheetz, 2006] and in immunology in particular [Huse, 2017]. In the context of immunology, T cells, and the T cell receptors (TCRs), have a special significance in being the very first players in adaptive immunity. Mechanics of T cells has been studied using a variety of techniques [Saitakis et al., 2017], recently revealing that T cells have atypical mechanical responses [Wahl et al., 2019, Yuan et al., 2021]. Likewise, mechanics of the interaction of the TCR and its molecular partner, the peptide loaded Major Histocompatibility Complex (pMHC), is a subject of current research with some groups reporting a catch bond [Kim et al., 2009, Liu et al., 2014], and some others not [Limozin et al., 2019]. A key to understanding how molecular scale mechanics and chemical kinetics are translated to cell scale mechanical behavior may be the bio-chemical link between intracellular moiety of molecular linkers and the cell cytoskeleton [Roy and Burkhardt, 2018, Blumenthal and Burkhardt, 2020]. The identity of the chain of proteins that form this link, often forming a molecular clutch, is well-known from experiments on non-immune cells, and for adhesion molecules like integrins where a hierarchy of actin-binding proteins like talin and vinculin, among others, are recruited to clusters of bound integrins [De Belly et al., 2022]; however, the nature of this link is still elusive for TCR where it has been called a condensate [Ditlev et al., 2019], perhaps to emphasize the physical, rather than chemical, nature of the interactions.

Cytoskeletal reorganizations are essential for correct functioning of leukocytes, including response after activation [Hivroz and Saitakis, 2016, Comrie and Burkhardt, 2016, Huse, 2017, Roy and Burkhardt, 2018, Puech and Bongrand, 2021, Göhring et al., 2022]. Like in other cell types, leukocytes, including T cells, exert forces mainly through their actin cytoskeleton. Forces are generated as a result of actin polymerization/branching and myosin-induced contractions. The details of rearrangement of the actin meshwork during adhesion and spreading was reported for T cells [Fritzsche et al., 2017, Ashdown et al., 2017]. The polymerization of actin at the cell edge leads to spreading [Bunnell et al., 2001, Dillard et al., 2014] and to actin retrograde flow close to the cell interface, that drags newly formed clusters of TCR towards the center of the spreading cell [Hartman et al., 2009]. This drag force, of frictional origin, to which all membrane receptors linked to the interfacial actin cytoskeleton - including both TCR and integrins - are exposed, is transmited through the linkers to the underlying substrate [Hartman et al., 2009, Dillard et al., 2014, Wahl et al., 2019], which in turn has been shown to lead to sustained signaling [Babich et al., 2012].

While the cross-talk of the cytoskeleton with signaling is well documented for T cells [Thauland et al., 2017, Colin-York et al., 2019], the details of the signaling cascade associated with mechanotransduction has been reported in only a few studies [Bashour et al., 2014, Hui et al., 2015, Wahl et al., 2019, Pathni et al., 2022]. It was shown that T cells can be activated simply by force application on TCR alone [Hivroz and Saitakis, 2016], via a Src kinase-dependent process [Li et al., 2010]. It is thus clear that force is an important control parameter of molecular function (especially in leukocytes). Interestingly, unlike in most other cell types, the sensing of mechanical environment in T cells appears to be myosin independent [Dillard et al., 2014, Wahl et al., 2019]; the extent of its spreading, when mediated by TCR alone, is biphasic with substrate stiffness [Wahl et al., 2019, Yuan et al., 2021]. T cells spread increasingly better on stiffer substrate, but only up to a point, after which the harder the substrate, the lesser the spreading [Judokusumo et al., 2012, O’Connor et al., 2012, Wahl et al., 2019, Yuan et al., 2021]. Such a behavior can be a result of the TCR-ligand bond being a catch bond, as modelled in the context of early spreading of fibroblasts [Oakes et al., 2018], but it could also be explained by a model that considers the mechanics and kinetics of the entire molecular assembly that links the cytoskeleton to the substrate [Wahl et al., 2019].

The role of the membrane-to-cortex attachment in regulating cell protrusions was recently emphasized for formation of cell protrusions in general [Welf et al., 2020]. In the context of integrin mediated adhesion, they can stabilize robust cell adhesion under flow [Whitfield et al., 2014], and mediate leukocyte rolling [Sundd et al., 2012]. Similar elongated membrane structure like microvilli play an essential role in the exploration of its environment by a T cell [Brodovitch et al., 2013, Cai et al., 2017], via TCR molecules located to the tip of the structure [Jung et al., 2016]. In all these examples, the link between receptors and cytoskeleton is difficult to characterize mechanically due to access issues.

The application of a force localized to the membrane achieved eg. by using an antibody as a molecular handle allows to test the links to the extracellular part of a specific membrane-bound receptor [Evans et al., 2005, Heinrich et al., 2005]. Pulling on these links, for example to break a ligand/receptor bond, may eventually link to extruding thin membrane tubes, and is one of the popular methods to probe membrane tension and mechanics [Hochmuth et al., 1996].

Tubes can be extracted from pure membrane systems such as giant unilamellar vesicles (GUV) [Dasgupta and Dimova, 2014, Bo and Waugh, 1989] or even artificial membranes [Dols-Perez et al., 2019]. Theoretically, the extrusion of nanotubes has been studied via analytical models [Derényi et al., 2002] and Monte Carlo simulations [Koster et al., 2005]. This models allow to link the force-extension curve of the tube’s extrusion to the mechanical properties of the membrane. Even if a small force overshoot can be seen, the experimental force vs. time curves of GUVs tube pulling are essentially monotonous [Derényi et al., 2002, Nowak and Chou, 2010].

Moreover, tubes can also be pulled from living cells [Borghi and Brochard-Wyart, 2007, Campillo et al., 2013], in order to probe viscoelasticity of the cell [Nawaz et al., 2012, Lu and Anvari, 2020], and can be complicated by the presence of the membrane to cytoskeleton links under force [Evans et al., 2005, Afrin and Ikai, 2006, Diz-Muñoz et al., 2010, Paraschiv et al., 2021]. Such experiments are usually described theoretically via models that take into account the viscoelasticity of the cell [Derényi et al., 2002, Lim et al., 2006, Brochard-Wyart et al., 2006, Schmitz et al., 2008, Al-Izzi et al., 2020], including in the context of deadhesion from the cytoskeleton [Nowak and Chou, 2010, Evans et al., 2005, Schmitz et al., 2008]. The theoretical analysis is complicated by the need to take into account the presence of membrane-to-cortex attachment (MCA) molecules [Sitarska and Diz-Muñoz, 2020]. The (few) existing theoretical studies are almost limited to numerical analysis [Paraschiv et al., 2021], which allow a comparison with extrusion experiments but do not permits a direct fit of the extrusion curve. Of note, experimental curves very often show a “peak then plateau” shape [Diz-Muñoz et al., 2010, Bretou et al., 2014, Sadoun and Puech, 2017], and only the plateau force is used to estimate the membrane tension and/or an attachment energy to the cytoskeleton, not providing any details of a molecular mechanism between the probed protein(s) and actin, but rather global membrane/cortex attachment [Krieg et al., 2008, Diz-Muñoz et al., 2010]. The precise significance of this peak, that is a reminder of the breaking of a bond, has been rarely adressed both experimentally and theoretically [Nowak and Chou, 2010].

An integrated mechanical model including effects of membrane, actin cortex and specific receptors is so far missing. In the present work, we propose a contribution under the form of an analytical model that allows to fit the force-elongation curves of nanotube extrusion, and considers explicitely the presence of the force peak upon retraction, before a plateau-like regime. We also describe in the same model the case where this peak is absent. Although the model does not give access to the underlying molecular mechanisms of the extrusion (i.e. phase transition of the phospholipids), it permits to extract separately the effective contributions of the elasticity provided by the membrane and the molecules. We demonstrate its efficiency by studying the case of the proteins composing the immune synapse, probed at the membrane of a living T cell. These proteins were predicted to exhibit differential interaction strengths with the actin, allowing the apparition of complex, biphasic, spreading behavior on activating substrates [Wahl et al., 2019]. The link of these proteins to the actin cortex represents an essential mechanism linking molecular structures, such as the TCR and the adhesion molecules, to the mechanosensitive elements that participate actively in the early T cell activation [Puech and Bongrand, 2021].

Here we access the mechanics of the putative link between the main lymphocyte membrane receptors, among them the TCR, and the actin cytoskeleton by pulling membrane nano-tubes from T cells, using antibody-coated beads in an optical trap. The time evolution of the force is fitted using a viscoelastic model that consists of springs representing either molecular or cellular elasticity and dash-pots that take into account the cellular and tube viscosity. By analysis data using scenarios corresponding to cases where the membrane receptor detaches or not from the cytoskeleton during tube formation, we are able to separate cellular and molecular elasticity. Finally, we compared hundreds of curves from experiments using different antibodies as molecular handles to access various membrane bound receptors.

## Results and discussion

### Experimental system

To dissect the interaction between immune receptors and actin cytoskeleton, we used optical tweezers to pull membrane tubes from Jurkat T cells. The cells, non activated and gently adhered onto polylysine glass substrates, were used to contact, for short duration (≤ 1 sec) and weak pushing forces (≤ 20 pN), beads decorated with antibodies directed specifically against a given molecule (Fig. 1A, B), eventually leading to a small fraction of the events (≤ 30 %) corresponding to the pulling of membrane tubes (Fig. 1C, similar to earlier reports [Diz-Muñoz et al., 2010]) and leading to force vs. time curves of specific morphologies (Fig. 1D). To exploit the richness of these curves, we developped a mechanical model encompassing molecular and cellular scales, together with the dynamics of the tube pulling (Fig. 1E, see below).

**Figure 1:**
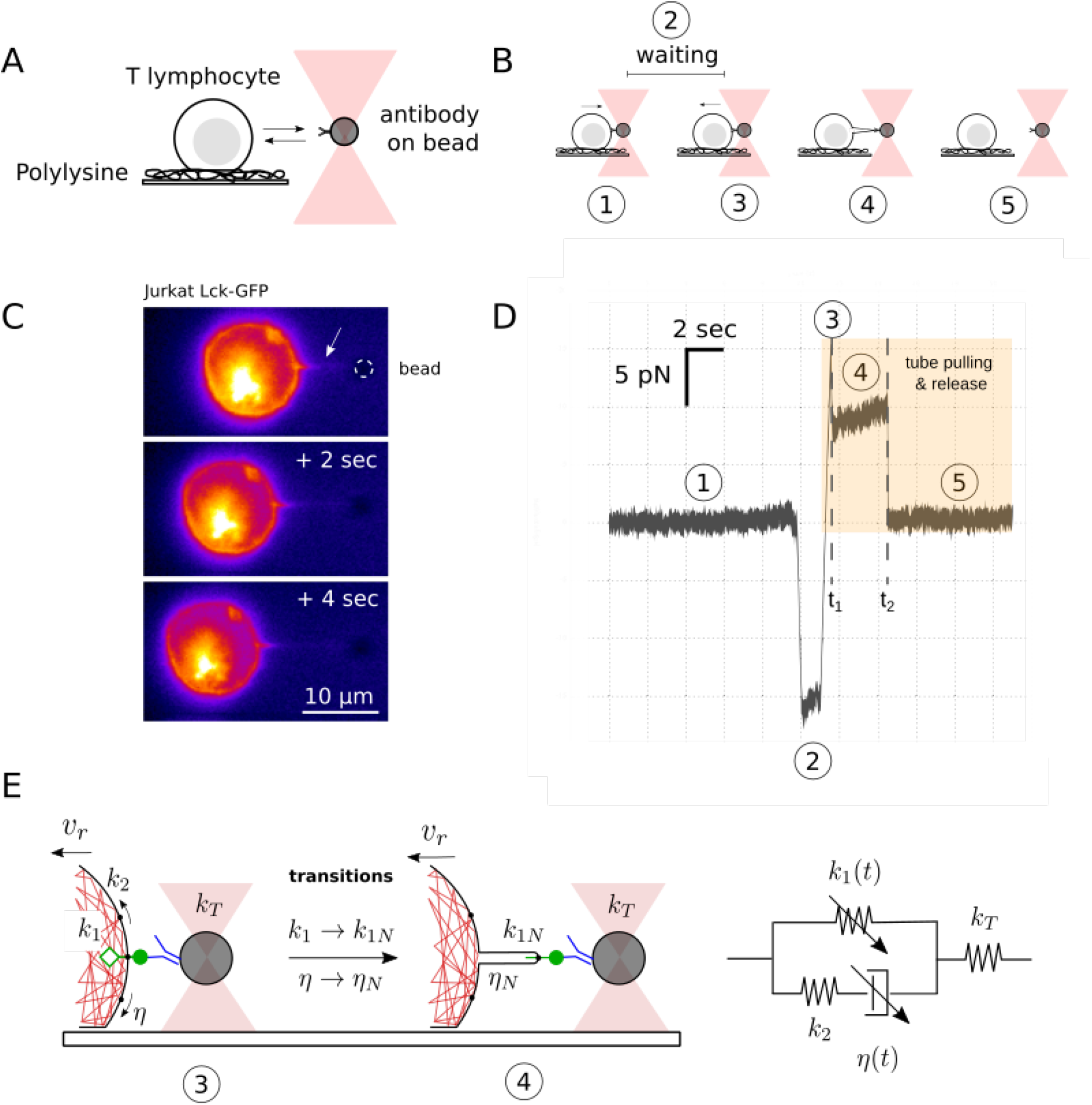
Schematics of experiments, example data and mechanical model of the OT-receptor-cell coupled system. A: The experimental setup consists of a colloidal bead coated with antibodies trapped using OT and a T cell adhered to a polylysine coated glass slide. B: The cell is put in contact with the bead ①, for a given duration ②, and then pulled back ③, eventually leading to the formation of a membrane tube ④, which eventually breaks ⑤. The time at transition from ③ to ④ is *t*_1_ and from ④ to ⑤ is *t*_2_. Steps ③ and ④ may be missing in some pulling cycles if the receptor-antibody bond breaks without a tube being pulled. C: Fluorescence micrographs of the process of tube pulling demonstrated in a membrane labelled T cell. D: Force vs. time curve during tube-pulling (labels correspond to stages shown in B). E: Details of molecular processes of interest and corresponding viscoelastic model consisting of a spring *k*_1_(*t*) representing the stiffness of the receptor-to-cytoskeleton link, in parallel with a series consisting of a second spring *k*_2_ and a dash-pot with viscosity *η* representing the effective rigidity and viscosity of the cell. A spring *k_T_*, in series with the whole, accounts for the stiffness of the optical trap. Note that *k*_1_(*t*) and *η*(*t*) are time dependant piece-wise functions that encompass the molecular and cellular transitions leading to the formation of a membrane tube.

To interrogate some of the essential transmembrane proteins involved in T cell activation [Limozin and Puech, 2019 Wahl et al., 2019], and also in IS formation, we used four molecular handles under the form of antibodies, to target the TCR/CD3 complex, the integrin LFA1 in its closed or open conformations and the long CD45 molecule (Fig. 3A). As positive and negative controls for the interaction with the cytoskeleton, we used that opened LFA1 is known to have a stronger interaction with actin than its closed or intermediate conformation [Limozin and Puech, 2019]. To our knowledge, the situation is largely unknown for the TCR/CD3 complex [Limozin and Puech, 2019, Wahl et al., 2019], and no clear data exists for CD45 [Cairo et al., 2010]. To destabilize the actin cytoskeleton, hence perturbating its possible links to the probed molecules, cells were challenged with a low concentration of Latrunculin A (hereafter LatA).

**Figure 2:**
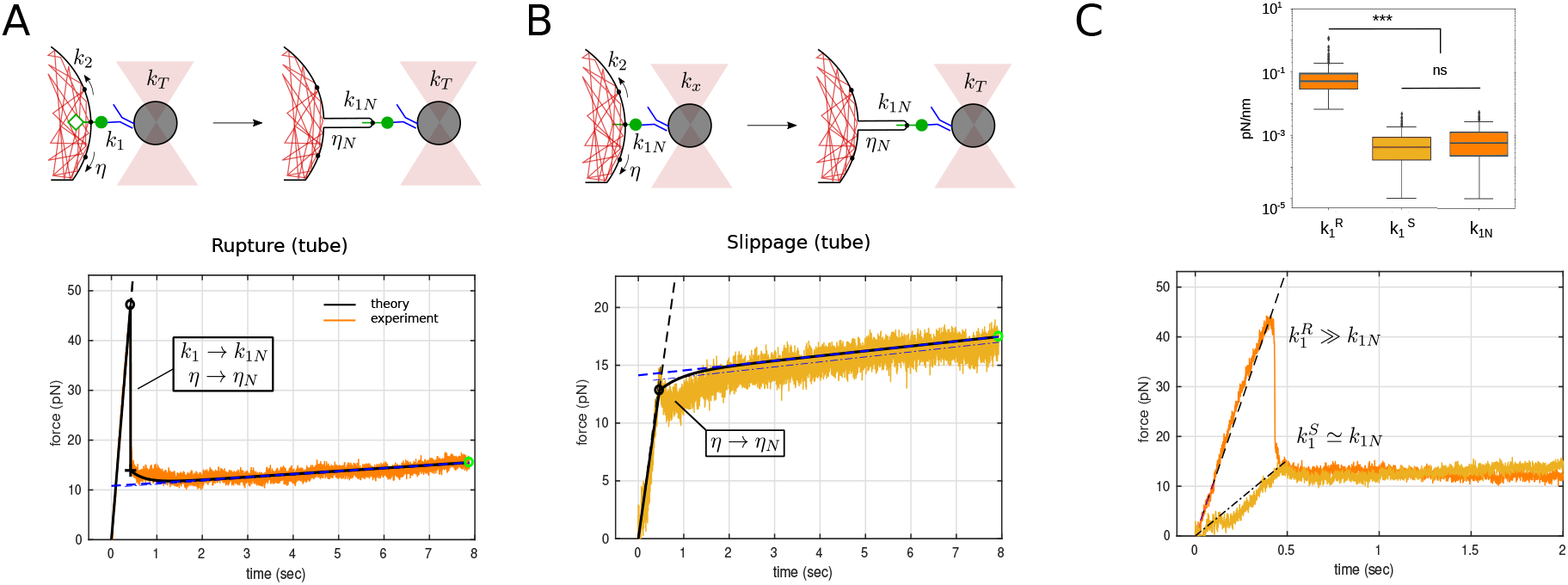
Schematic of microscopic events and corresponding data. A. Top: rupture of the receptorcytoskeleton link (hollow green diamond) leads to tube formation, with corresponding changes in viscoelastic parameters. Bottom: corresponding force curve showing a discontinuous jump upon rupture B. Top: in absence of a receptor-cytoskeleton link, a tube is pulled simply with membrane “slippage” on the cytoskeleton, implying a transition only of *η*(*t*). Bottom: the force curve shows a simple discontinuity and no jump. C. Top: *k*_1_ and *k*_1*N*_ for rupture (R, N=116 for each) and *k*_1_ for slippage (S, N=165) events. Bottom: overlay of typical rupture and slippage force curves to graphically emphasize that, after the rupture event at *t*_1_, the two cases are identical.

**Figure 3:**
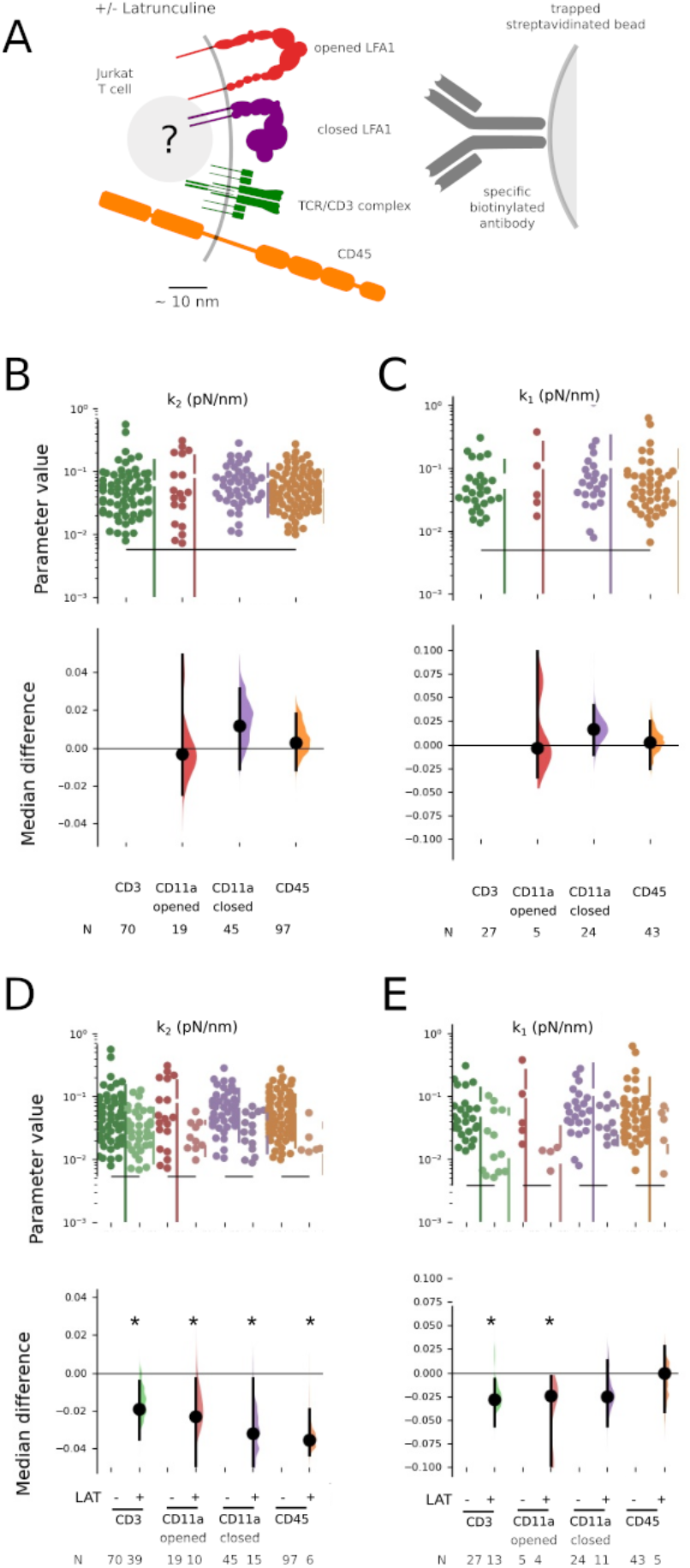
Varying the antibody handle or perturbing the cytoskeleton. A. Schematic of the receptors and their conformations which were specifically solicited by appropriate antibodies on the bead used to pull membrane tubes. B. Estimation plots (from Dabest, [Ho et al., 2019]) of *k*_2_ (effective cell stiffness) taking CD3 as a reference. No difference between the handles is seen. C. Same as B, for *k*_1_ (stiffness of receptor-cytoskeleton link). Again, no difference between the handles is seen. D. Comparing *k*_2_ before and after disruption of cytoskeleton using Lat A. In each case, *k*_2_ decreases, coherent with a global mechanical perturbation of the cell when the actin is perturbed. E. Same as D but for *k*_1_. Differences emerge after treatment with LatA for CD3 and CD11a open cases, indicating that an interaction exists between the receptor and the actin cytoskeleton, unlike for CD11a closed and CD45 ones. One dot corresponds to one fitted curve. The corresponding values of *k*_1*N*_, *η* and *η_N_* can be found in Fig. S5. N indicates the number of curves for each case. Star (*) indicates significant difference of medians following Dabest analysis (see SI).

### Force curves morphologies and transitions

Visual inspection of roughly 8900 curves revealed four morphologies (Fig. S1). First, and most interesting, about 4% of the curves exhibit a clear spike-like discontinuity followed by a slow increase and a second discontinuity where the antibody-receptor bond breaks and the force goes to zero, henceforth called “rupture” case Fig. S1A). Second, 6% show a step-like discontinuity followed by slow increase and a step down to zero force, henceforth called “slippage” (Fig. S1B). Third, 23% exhibit a spike which immediately falls to zero force, called “detachment” (Fig. S1). As expected, due to short and gentle contact parameters imposed in order to fulfill single molecule conditions, a fourth case is seen in the vast majority (67%) of the curves, where no attachment of the bead to the cell occurs, and no meaningful force-curve is obtained (denominated “contact”). Of note, the slowly rising plateau seen in the first two cases is characteristic of tube extraction [Evans et al., 2005, Schmitz et al., 2008, Diz-Muñoz et al., 2010].

We interpret the difference between the two tube cases in molecular terms. In the rupture case, the spike/discontinuity corresponds to the rupture of the cytoskeleton-receptor link and a concomitant tube formation, which were not experimentally separable in time (Fig. S1A). In the slippage case, the receptor-to-cytoskeleton link is either absent or very weak, and the membrane slips over the actin cortex and a tube forms without having to rupture any specific linkage (Fig. S1B). Finally, the force abruptly falling to zero, seen in the detachment case, and eventually at late times for tubes, corresponds to the breaking of the extracellular antibody-receptor bond, leading to the detachment the bead from the receptor handle. In some cases, the tube was not rupturing at the end of the experiment, due to a finite total pulling length hence duration, leading to “infinite” tubes. All these cases can be interpreted in the frame of our mechanical model.

### Mechanical model

The relevant part of the experimental system and its equivalent mechanical model are pictured in Fig. 1E. The mechanical model is essentially a standard linear solid model [Schmitz et al., 2008, Lim et al., 2006] representing the cell-tube-receptor system, in series with another spring to account for the optical trap. The former consists of a spring with spring constant *k*_1_(*t*) that represents the stiffness of the receptor-to-cytoskeleton link as well as the tube that is to be pulled, in parallel with a second spring, *k*_2_, and a dashpot, with viscosity *η*(*t*), representing the effective rigidity and viscosity of the cell. An additional spring *k_T_*, in series with the whole, accounts for the stiffness of the optical trap. It is important to include *k_T_* as it was previously shown that neglecting the stiffness of the handle - here the OT - may lead to significant over or underestimation of the mechanical properties of molecules [Manca et al., 2013, Bellino et al., 2019].

At time *t* = *t*_1_, the receptor-to-cytoskeleton link ruptures and the membrane detaches from the cytoskeleton leading to the formation of the tube. The stiffness of the link (*k*_1_) is not expected to be time dependent while it is intact, and similarly, the stiffness of the tube (*k*_1*N*_) is considered to be time independent. The cell elasticity (*k*_2_) is not expected to be impacted by tube pulling, however, the viscosity, with potentially major contribution from the membrane itself, may change (from *η* to *η_N_*). Thus, *k*_2_ is constant and *k*_1_(*t*) and *η*(*t*) are piece-wise constant. *k_T_* is experimentally set and constant while the tube exists. At the end, the tube detaches due to deadhesion of the receptor-ligand bond, *k_T_* then (effectively) goes to zero and the force falls to the baseline value. This sequence is clearly reflected in the force curves (see example in Fig. 1D).

The constitutive model of the coupled system is then given by the following differential equation

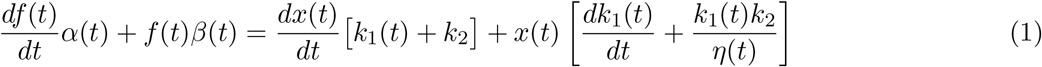

where 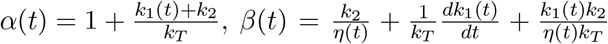. The imposed distance as a function of time is given by *x*(*t*) = *v_r_*(*t*) × *H*(*t*), where *H*(*t*) is a Heaviside function, *v_r_* is imposed at time *t* = 0 (which corresponds to *f* = 0 when starting to pull on the system, Fig. S1).

The response is evaluated by solving the differential constitutive equation separately before and after the discontinuity at *t* = *t*_1_. The analytical solution, and its comparison with the numerical solution, can be found in SI (Eq. 2). This solution is a general case of the classical standard-linear-solid model (SLSM) [Schmitz et al., 2008, Lim et al., 2006], with an additional spring *k_T_*, and where time discontinuities are introduced for both *k*_1_(*t*) and *η*(*t*). The solution at *t* ≤ *t*_1_ deviates from a linear behavior expected from purely elastic contributions (*k*_1_ + *k*_2_), and describes the relaxation caused by the viscosity of the cell, *η*(*t*). The solution at *t* > *t*_1_ describe the relaxation of the system after the rupture of the link (*k*_1_ → *k*_1*N*_), and the concomitant transformation of the locally flat cell membrane into a tube (*η* → *η_N_*), which results in a plateau-like shape in the force evolution (Fig. S1 A). SI Eq. 2 was used to fit all the experimental curves in order to obtain the value of the mechanical parameters.

### Curve fitting and extracted parameters

The pipeline for fitting the curves consists of the following steps (detailed in SI). The raw force curves are smoothed, and categories (<u>R</u>upture tube, <u>S</u>lippage tube, <u>D</u>etachment, Contact) are determined by looking for discontinuities and extrema using in-build Matlab routines. Curve fitting range is also determined at the same time and constraints are chosen depending on categories.

Fitting is done on the entire chosen range such that each piece is fitted with at most 3 parameters, with range of parameters fixed according to SI Table 2. *v_r_* and *k_T_* are fixed experimentally, *t*_1_ is determined by direct detection of the discontinuity of the force-curves and the parameters *k*_1_, *k*_1*N*_, *k*_2_, *η*, and η_N_ are determined from the fit. In case of detachment, *k*_1*N*_ and *η_N_* do not exist since no tube is pulled. In case of slippage, *k*_1_ = *k*_1*N*_ is imposed since, in absence of the receptor to cytoskeleton link, there is no transition from pulling on the link to pulling on the tether. The rounded median values of the fixed and fitted parameters, pooling data from all conditions, are given in Table 1.

**Table 1:** Physical parameters of the model (median values on the entire data set). R: Rupture, tube. S: Slippage, tube. D: Detachment, no tube. Green / red symbol: parameter accessed or not by the model (resp.)

| Parameter | Symbol | Value | Units | R | S | D |
| --- | --- | --- | --- | --- | --- | --- |
| Discontinuity time | $t_1$ | 0.25 | s | ✓ | ✓ | ✗ |
| Molec. stiffness | $k_1$ | 0.05 | pN nm<sup>-1</sup> | ✓ | $k_{1N}$ | ✓ |
| Tube stiffness | $k_{1N}$ | 0.0005 | pN nm<sup>-1</sup> | ✓ | ✓ | ✗ |
| Cell stiffness | $k_2$ | 0.05 | pN nm<sup>-1</sup> | ✓ | ✓ | ✓ |
| Cell viscosity | $\eta$ | 0.04 | pN nm<sup>-1</sup> s | ✓ | ✓ | ✓ |
| Tube viscosity | $\eta_N$ | 0.008 | pN nm<sup>-1</sup> s | ✓ | ✓ | ✗ |
| Pulling velocity | $v_r$ | 2500 | nm s<sup>-1</sup> | - | - | - |
| Trap stiffness | $k_T$ | 0.25 | pN nm<sup>-1</sup> | - | - | - |

While *k*_1_ is explicitly determined here for the first time, the obtained values of other mechanical constants are overall coherent with literature [Evans et al., 2005, Schmitz et al., 2008]. Explicitly, Ref. [Evans et al., 2005] reported a value equivalent to *k*_1_ + *k*_2_ = 0.3 pN/nm which compares well with our value of 0.1 pN/nm for *k*_1_ and *k*_2_; Ref. [Schmitz et al., 2008] reported values equivalent to *k*_2_ = 0.2 pN/nm (0.05 pN/nm here) and *k*_1*N*_ = 0.001 pN/nm (0.0005 pN/nm here) (see Table 1). The model turns out to be robust for *k*_1_, *k*_1*N*_ and *k*_2_, but much less for *η*, whose obtained values are widely dispersed. Parametric study (see below) reveals that the fit is not very sensitive to *η*. Nethertheless, its values are coherent with litterature [Schmitz et al., 2008].

### Mechanical transitions observed between the different tube morphologies are coherent

On one hand, as prescribed by our fitting to the model, *k*_1_ > *k*_1*N*_ in the “rupture” case. On the other hand, we observe that the values for *k*_1*N*_ are similar for the “rupture” or “slippage” cases for tubes (Fig. S1C and S6A,B), corresponding to the fact that the things become similar when the intracellular bond is broken and *k*_1_ reaching *k*_1*N*_ (“rupture”) and when starting from it (“slippage”). Interestingly, *k*_1_ is similar for detachment events and “rupture” tubes (Fig. S6A), while *k*_2_ is not dependent on the event being a tube or a detachment (Fig. S6C). Moreover, the viscosity *η_N_*, ie. after all transition(s), is the same for the two cases with a tube, corresponding to a similar tube pulling mechanism. Importantly, all of these observations are independent from the precise molecular handle that was used to pull the tubes, showing the consistency of our model and methodology. Interestingly, one can appreciate that *k*_1*N*_ is not affected by LatA, while *η* seems to be decreased in all cases, together with *η_N_* (Fig. S6).

We present the distribution of times *t*_1_ and *t*_2_ on Fig. S7A,B without and with LatA, respectively. *t*_1_ corresponds to the time of the first transition. In the rupture case, it is the simultaneous transition of *k*_1_ and *η*, while for slippage case, it is the transition of *η* alone. Fig. S7C shows no difference of *t*_1_ between rupture and slippage cases. This validates our approximation that the two transitions are detected simultaneously for the rupture case in our experiments.

### Immune receptor interactions with cytoskeleton are molecule specific

Fig. 3B, C present the cellular elasticity, *k*_2_, and the molecular bond parameter, *k*_1_, which correspond to the intracellular bond of the handle to the cytoskeleton, respectively. The results obtained for *k*_1*N*_, *η* and *η_N_* can be found on Fig. S5. None of the five parameter appears to be affected by the particular handle used, which allows to conclude that the molecular details of the *extracellular* interaction between bead and cell are not affecting our measurements.

Notably, the low doses of LatA that were used affected the global cell mechanics, as expected, which can be seen on their homogeneous and significant effect on *k*_2_ values (Fig 3D). Remarkably, LatA did not affect the intracellular molecular bond parameter *k*_1_ the same way for the different handles (Fig 3E). While a strong and significant effect is seen for the opened conformation of LFA1, no significant effect can be seen for the closed conformation, even if the median shift is similar, in agreement with the relative interactions of the two conformations with actin. Interestingly, the case of CD45 was not showing any sensitivity to the drug. For TCR/CD3, we observed a significant effect of the drug. Taken together, we see a differential effect of the drug on *k*_1_ that we interpret as a differential interaction with the cytoskeleton. Let us now explore in greater detail the possible meaning of *k*_1_, that, by its very nature bridges the molecular scale (a few nm), and the mesoscopic tube scale (100nm).

To do so, *k*_1_ is further decomposed into a circuit of springs as shown in Fig. S8, such that 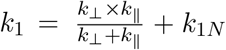. From table 1, we already know that 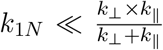. We will further require that in an unperturbed state, *k*_∥_ < *k*_⊥_: a statement that we will substantiate below.

Physically, we identify *k*_⊥_ as the elasticity of the direct link between the intracellular part of the surface-molecule that is bound to the antibody handle and which transmits a force locally roughly perpendicular to the membrane and parallel to its own mechanical axis. *k*_⊥_ is a local property of the solicited receptor and is therefore expected to be antibody dependent. *k*_∥_ represents the elasticity associated with the weaker links of other membrane bound receptors which are necessarily pulled along when the membrane is pulled, a mechanism somewhat similar to the frictional breaking proposed by Groves [Yu et al., 2013] or stick-slip mechanisms mooted in the context of mechanosensing [Wahl et al., 2019], with the force being transmitted parallel to the membrane. *k*_∥_ is a non-local, mesoscopic parameter, which is not specific to the receptor that is bound and therefore it is expected to be antibody independent. *k*_1*N*_ can be thought of as the elasticity associated with the emerging tube when it comes into being, and probably represents residual non-specific interaction between the bulky intracellular moieties of membrane-bound receptors and the intracellular environment, making it the softest spring in the circuit.

This relatively simple mechanical circuit captures the behaviour of all the antibodies tested under force and in presence/absence of latrunculin. Let us consider each case separately.

- *The unperturbed system*: for all four antibodies, namely aCD3, aCD11a-open, aCD11a-closed and aCD45, the shape of the circuit ensures that the response is dominated by the softer spring in series - namely *k*_∥_. *k*_⊥_, which is expected to be the antibody dependent element but also the stiffest element in the circuit, is not probed. Naturally, whatever the antibody, the response is identical, dominated by the non-specific *k*_∥_ component, measured here to be 0.05 pN/nm. The only parameter reported in literature that is akin to *k*_⊥_, was for CD3, in the context of cell spreading and mechnaotrasduction [Wahl et al., 2019], where it was reported to be 0.3 pN/nm - an order of magnitude stiffer than *k*_∥_ reported here - consistent with our hypothesis.
- *Perturbation of the actin cytoskeleton*: perturbing actin using latunculin is expected to strongly impact the direct link between the target receptor and the actin cytoskeleton. With a large dose of latrunculin, we can expect all existing linksto be severed. However, at the small dose of latrunculin used here, we are left with many cases of rupture, indicating that in these cases, the link survives and it is these cases where we measure *k*_1_. Latrunculin, at low doses, should not impact *k*_∥_, which is expected to depend on the meso-scale connectivity of the actin network.

In all cases, at high force or extension, the link, presumably *k*_⊥_, ruptures and the response is now dominated by *k*_1*N*_.

It is important to note that the attribution of each element to specific molecular players is, given the state of art, necessarily speculative but the basic mechanical arguments based on the spring network is strictly validated by our experimental data.

### Model exploration and predictions

To assess the robustness of parameter determination, we performed a parametric study of the model (Fig. 4), to dissect the effects of variations of the different fitting and fixed parameters. As expected, the early-time quasi-linear behavior is mainly governed by *k*_1_, which does not affect the post-rupture part of the curve (Fig. 4A). To the contrary, the value of *k*_1*N*_ affects only the residual slope of the force for *t* > *t*_1_ (Fig. 4B). Coherently with our observations made when fitting the data, variations in tube viscosity *η* has minimal impact before *t*_1_, and only a moderate one after, (Fig. 4C). Aside, *η_N_* governs the trend of the force from a convex to a concave behavior for *t* > *t*_1_ (Fig. 4D). Aside, the shape of the relaxation (concave or convex) depends on the value of *t*_1_ (Fig. S9). Interestingly, large variations of *k*_2_ have only a small impact on the linear loading phase, but *k*_2_ however plays a crucial role for *t* > *t*_1_ (Fig. 4E), and controls for the slippage case the maximal force at *t*_1_ and curvature after it (Fig. S10). Notably, the behavior of the force-curve also depends on the stiffness of the force transducer, and we scan the typical range of common force-spectroscopy measurements, going from photon-field (softer) to mechanical (stiffer) transducers, showing the profound impact of the measuring spring on the morphology of the force vs. time data curve (Fig. 4F) [Bustamante et al., 2000].

**Figure 4:**
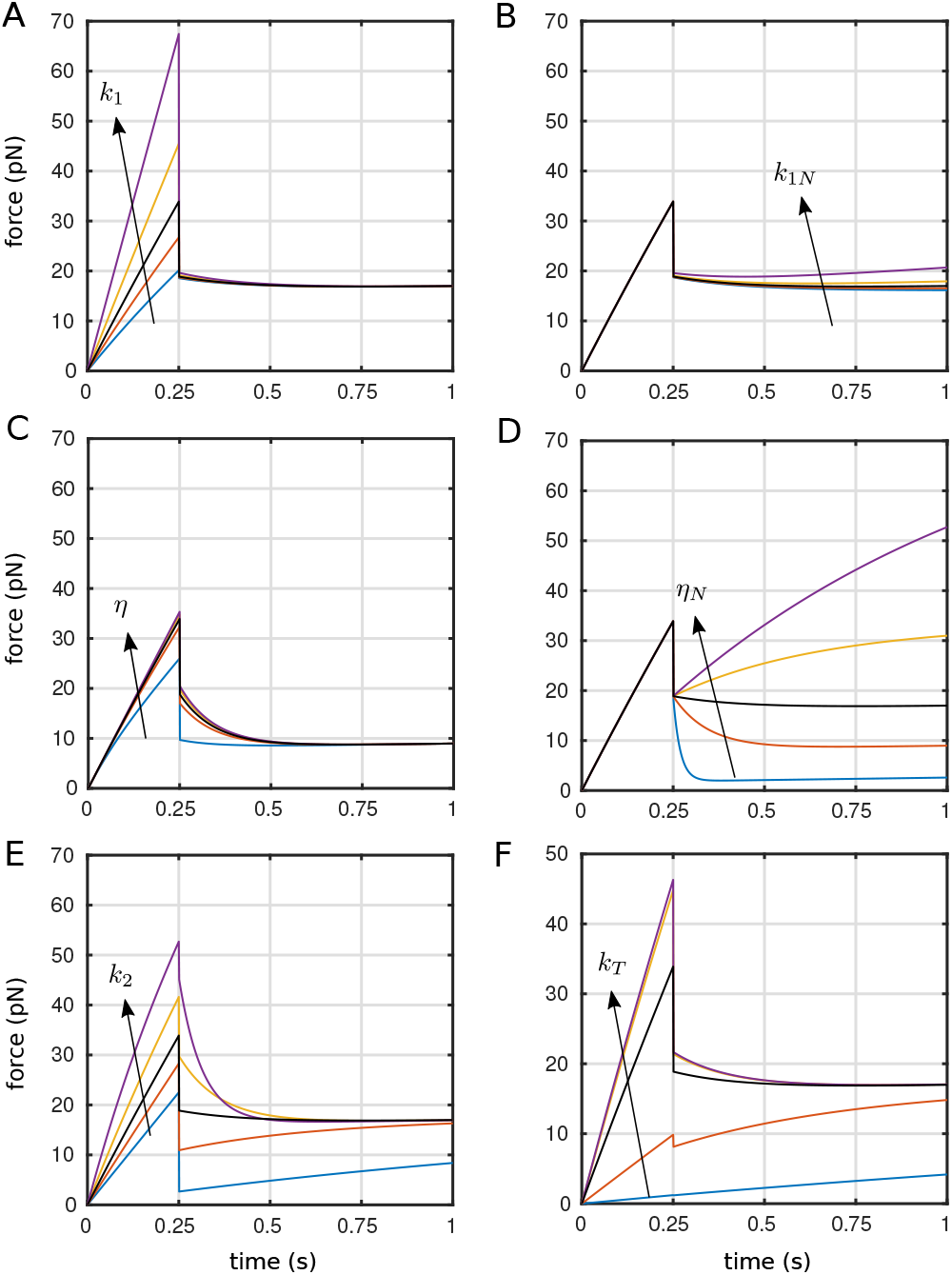
Parametric study of the viscoelastic model for the rupture case. The black curve inbetween the others correspond to the one obtained via the fitting in Fig. S1, left panel. For panels from A to F, parameters are *v_r_* = 2000nms^-1^, *k_T_* = 0.25pNnm^-1^, *k*_1_ = 0.05pNnm^-1^, *k*_2_ = 0.05pNnm^-1^, *η* = 0.04pN nm^-1^ s, *k*_1*N*_ = 0.0005pN nm^-1^, *η_N_* = 0.008pN nm^-1^ s, and *t*_1_ = 0.25s. The others curves have been obtained multipling these values by the following vector of factors {0.1, 0.5, 1, 2, 5}. For panels E and F, parameters are *k_T_* = 0.01, 0.1, 1, 10, 100 pN nm^-1^, and *t*_1_ = 0.5, 0.75, 1, 1.25, 1.5 s, respectively.

Overall, we explored a wide range for the values of the parameters, and conclude that the model’s predictions – both qualitative and quantitative – are robust. Most importantly, the model is highly sensitive to *k*_1_, which is the principle parameter of interest in the present study.

## Conclusions

Our data, model, and the fitting presented here demonstrate that pulling of a tube may (rupture case), or may not (slippage case), involve breaking of an internal bond, distinct from the external antibody/antigen bond. The signature of this breaking is contained in the force curve (fig. 1) where an increase followed by an abrupt fall signifies rupt whereas a gentle increase and smooth fall signifies slip. This interpretation is supported by previous work of Nowak et al who showed that pulling of cytoskeleton-associated membrane tubes involves higher force and a more abrupt jump as compare to pulling pure membrane tubes [Nowak and Chou, 2010]. Concentrating on the former (rupture) case, the internal rupture involves snapping of the spring k_1_ in Fig. 1, which upon rupture takes on its residual value of *k*_1*N*_. We have further shown that the value of *k*_1_ is latranculin dependent for some antibodies but not others. Our quantitative analysis of the cellular and molecular parameters of our model is then coherent with our precedent modelling of T cell bi-modal spreading [Wahl et al., 2019].

Overall, we can conclude that the intracellular molecular spring linking the receptor to the cytoskeleton, with stiffness k1, originates from two main components that are not explicitly introduced in our 1D model of dynamical tube pulling, in addition to its residual value (*k*_1*N*_) associated with the tube after bond rupture. These two main components are: *k*_⊥_ which is a spring-like element perpendicular to the local membrane plane and colinear to the traction force. It corresponds to the actual molecular interaction with the actin, and *k*_∥_, an element parallel to the membrane, encompassing the interaction of the cytoskeleton with the rest of the membrane including proteins which spans it. The first element is drug dependent and we propose that the second, which corresponds to an intermediate scale is essentially independent of the exact details of actin to receptor interaction (Fig. S8).

The interpretation presented above show that we do probe differential interactions of IS proteins with the actin cytoskeleton by using different antibodies as molecular handles, but the difference cannot be revealed without perturbing the system using a drug. The stiffness response of the molecular spring corresponding to the specific link to actin, *k*_⊥_ is “hidden” due to the presence of a softer spring in the system. We can only state that for all receptors targeted here, the value of *k*_⊥_ in an unperturbed cell is larger than 0.05 pN/nm. To dissect the exact values or origin of *k*_⊥_, more refined experiments, such as using cells with specific KOs of ERM molecules, talin or other putative adaptor molecules needs to be performed, with large enough data-sets to reveal potentially subtle differences in the fitted parameters. Nevertheless, here we demonstrated the importance of the mesoscale, represented by the membrane and its association with the actin network, for a full understanding of the IS at the molecular scale. A proper understanding of this intermediate scale may be key to how immune cells convert molecular cues to cell scale activation.

## Material and Methods

Details about experimental and numerical procedures can be found in the supplementry materials section.

## Funding and Acknowledgments

The project leading to this publication has received funding, as a postdoc grant to FM, from France 2030, the French Government program managed by the French National Research Agency (ANR-16-CONV-0001), from Excellence Initiative of Aix-Marseille University - A*MIDEX, and from Labex INFORM (ANR-11-LABX-0054) and A*MIDEX project (ANR-11-IDEX-0001-02). This work was supported by the GDR ImaBio through master’s internships funding (GE, ON). The authors thank M. Biarnes-Pelicot, the PCC facility and JPK Instruments/Bruker for continuous support, and A. Sunčana Smith for fruitful comments and discussions.

## Supplementary Material for Manca. et al.

### Content

- Material and Methods
- Supplementary Figures
- SI References

## Materials and methods

### Cell culture

Jurkat E6-1 cells (ATCC, #TIB-152) were grown at 37°C and 5% CO_2_ in Roswell Park Memorial Institute (RPMI) 1640 1x medium (Gibco, #11875) supplemented with 10% fetal bovine serum (FBS) and 1% stabilized L-glutamine (GlutaMAX, Gibco). They were diluted in fresh medium every 2 or 3 days in order to keep the concentration between 0.4 10^6^ cells/mL and 1.2 10^6^ cells/mL.

### Beads preparation

We used polystyrene beads of diameter 2 *μ*m pre-coated with streptavidin (Polysciences, Inc., #24160), having an initial concentration of 3 10^9^ beads/mL. They were diluted to 1/10 in Dulbecco’s Phosphate-Buffered Saline 1x (DPBS) with 1% (w/v) Bovine Serum Albumin (BSA, Sigma Aldricht) to reach a volume of 250 *μ*L. They were washed three times by centrifuging during 6 min at 6,700 g and replacing the supernatant with 250 *μ*L of DPBS/BSA 1%.

After the last wash, beads were resuspended in 100 *μ*L of DPBS/BSA 1% and a solution containing 10 *μ*L of biotinylated antibody at 0.5 mg/mL was added. We used biotinylated mouse IgG2aK monoclonal antibodies (all from eBioscience, Thermofisher, Biorad): anti-CD45RO (clone UCHL1), anti-CD3 (clone OKT3, clone UCHT1), anti-LFA1 (closed conformation, anti-CD11a clone 38; opened conformation anti-CD11a clone HI111). We used as an isotype control a non specific IgG2aK (clone eBM2a). Beads and antibodies were co-incubated 30 min at room temperature (RT) under stirring. Beads were then washed as previously and finally resuspended in 500 *μ*L DPBS/BSA 1%, to a final concentration of ~ 10^8^ beads / mL. The functionnalized beads were stored during maximum one month at 4°C.

### Sample preparation

Petri dishes having 35 mm diameter and 0.17 mm thick glass bottom (Fluorodish, WPI, #FD35-100) were incubated 30 min at RT with 2 mL of polylysine solution (Sigma-Aldrich, #P8920) diluted to 1/10 and washed three times with 2 mL DPBS 1x.

Approximatively 5 10^5^ cells were taken from the culture one day after splitting, and resuspended in pure RPMI after gentle centrifugation during 3 min at 400 g; they were then transferred to the Petri dish. They were incubated 30 min in culture conditions to allow them to adhere. Medium was then gently replaced by supplemented RPMI and 10 μL of beads solution corresponding to 1-1.5 10^6^ beads just before installing the sample on the heating microscope stage.

To perturb actin, latrunculin A (Sigma-Aldricht) was used at a low final concentration of 2 to 5 μM, incubated 30 min with cells before the experiment. Note that the experiments were performed in the presence of the drug. We verified experimentally that the presence of a small residual amount of DMSO in the experiment medium did not affect the measurements.

### Optical tweezers

The acquisition of force curves was performed with a Nanotracker 2 (JPK Instruments/Bruker) optical trapping device, equipped with a motorized/piezo stage, mounted on an inverted microscope (Axio Observer, Zeiss). The sample was fixed on a thermoregulated petridish holder (PetriDish Heater, JPK Instruments/Bruker), the temperature of which was set to 37°C for all the experiments.

The trapping objective (C-Apochromat 63x/1.2 W Corr, Zeiss) was covered by a drop of immersion oil (Immersol W 2010, Zeiss) that has a refractive index near to the one of water (n=1.334 at 23°C). The detection objective (W-Plan-Apochromat 63x/1.2 W Corr, Zeiss) was immersed in the sample medium. The optical trapping laser had a wavelength of 1064 nm and a maximal power of 3 W. The laser was focused in the medium by the trapping objective and the out-coming beam was driven through the detection objective to quadrant photodiodes. These allow to measure the displacement of the trapped object in the back focal plane in three dimensions and to quantify the forces after calibration.

For transmission light microscopy, a LED lamp is focused on the sample by the detection objective and the picture is acquired by a CCD camera (DFK 31BF03.H, Imaging Source).

The NanoTracker software (version 6+ on GNU/linux, JPK Instruments/Bruker) controls the position of the objectives, the position of the sample, the position of the trap, the intensity of the laser and the attenuation filters before the detection photodiode.

The distribution of bead diameters was measured separately on bright field microscopy images and the average value was used in all experiments (2R ≃ 1.67 ± 0.07 *μ*m). We imposed a medium viscosity *η* of 6.96 10^-3^ Pa/sec. The stiffness of the trap is calibrated by the software based on the spectral analysis of the thermal noise implemented in the control software (1).

A ramp designer allows to program the motion of the sample with the piezoelectric stage. The ramps had three phases: first, a rectilinear motion toward the cell interrupted when the force detected by the photodiode exceeds a given threshold (10 or 15 pN); then, a pause of a given duration in which the sample stays immobile (0 to 1 sec) and the force relaxes; and finally, a rectilinear motion in the opposite direction until a given distance is reached (15-20 *μ*m). The speed of the forward and backward motions was typically set at either 2 or 2.5 *μ*m/sec. The acquisition frequency for the force curve data was 2048 Hz.

During the experiment, the force signal in three dimensions, based on the stiffness calibration along the three motion axis, is recorded and saved. In order to optimize the force to be colinear to the relative motion of the bead and cell, we attempted to have the trajectory perpendicular to the cell membrane. For this, to minimize lateral forces, when the trajectory was in X (resp. Y) axis we incrementally adjusted the Y (resp. X) and Z positions to minimize the force measured in Y (resp. X) and Z axis before the first cell / bead contact.

The measurements, which are saved as compressed and encoded commercial format files, were finally converted by using the NanoTracker data processing software (JPK/Bruker) into tab separated text files that can be feeded into our Matlab procedures.

### Model

The general solution of Eq. 1 (see main text) is given by

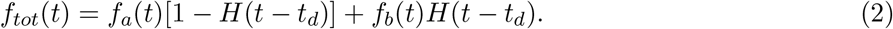

with

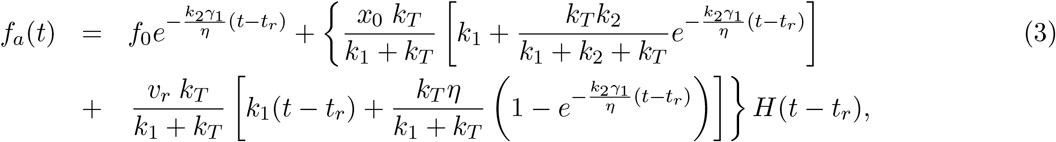

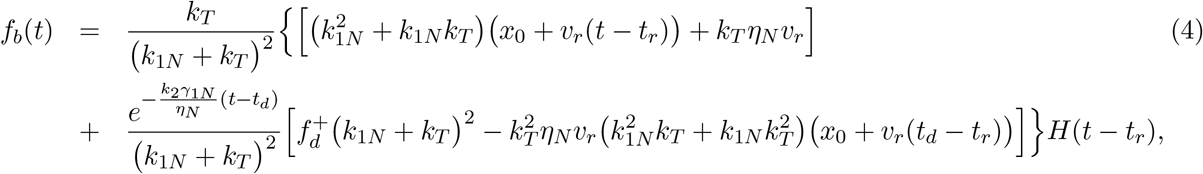

where, we considered *k*_1_(*t*) and *η*(*t*) as piecewise functions

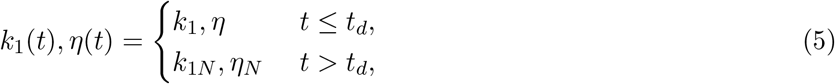

being *t_d_* the time at which the discontinuity happens, *t_r_* the retraction time ie. the time at which the retraction starts, *x*_0_ the initial position of the optical bead, *f*_0_ the initial force measured by the tweezers, *v_r_* the speed of pulling and 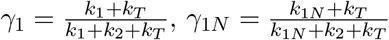.

The value of the force after the discontinuity is equal to

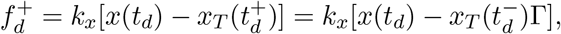

being *x*(*t*) the total length of the system, *x_T_*(*t*) the distance of the optical trap from its equilibrium position and 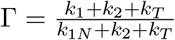 the ratio of the effective stiffnesses in *x_T_* before and after *t_d_*.

Finally, getting the recorded position of the optical bead from Eq. 2, 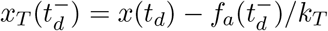, and being *x*(*t_d_*) = *v_r_* × (*t_d_* – *t_r_*), we have

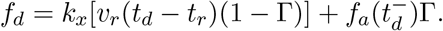

Notice that, with *t_d_* = *t*_1_, and explicitating the force 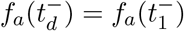, we get the explicit solution of *f_b_*(*t*) (see Eq.7), which is a function of the free parameters *k*_1_, *k*_1*N*_, *k*_2_, *η*, *η_N_*, *t*_1_ only.

From Eq.7, with the boundary conditions we choose to offset the raw data to, which are *f*_0_ = 0, *x*_0_ = 0 and *t_r_* = 0, we obtain the following simplified form of the time-force evolution

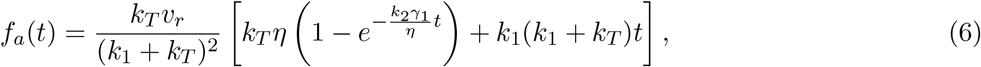

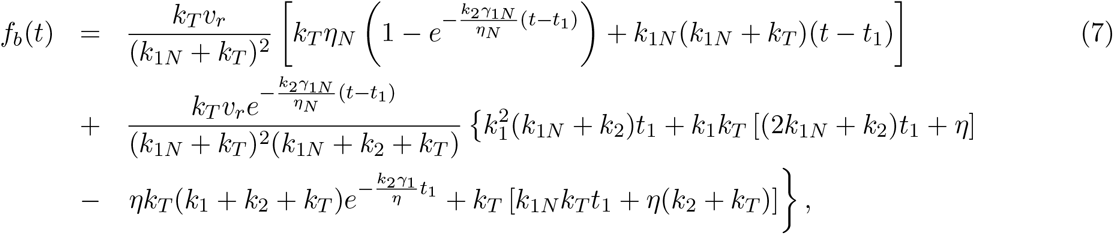

Notice that, in Eq.6, for *k_T_* → +∞, *k*_1_/*k_T_* → 0, we recover the classical solution of the standard-linear-solid model (SLSM) (2,3). With Eq. 2, we fitted all the experimental curves and obtained the distribution of the parameters.

In Fig. S2, we report the theoretical results, both numerical and analytical, of the force evolution *f_tot_*(*t*) obtained for the input *x*(*t*). The system parameters have been fixed to values similar to the ones reported in Table 1: *k_T_* = 0.25pN nm^-1^, *k*_1_ = *k*_2_ = 0.05pNnm^-1^, *η* = 0.02pN nm^-1^ s, *k*_1*N*_ = 0.0005pN nm^-1^, *η_N_* = 0.004pNnm^-1^ s, and *t*_1_ = 0.25s. The red curve corresponds to the numerical resolution of Eq. 1 (with a Runge Kutta 4th order integrator) and the dotted-black curve corresponds to the analytical solution of the force of Eq. 2. For reasons of numerical stability, the Heaviside functions *H*(*t*) have been replaced by a step-like function defined as: 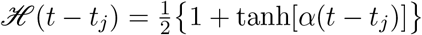, with *α* = 10^7^.

### Data Analysis

The analysis of the force measurements consisted of three major steps. First, the classification of the curves, which automatically identifies their characteristics. Second, the fitting of the curves, which gives the estimation of the mechanical parameters. Third, the statistical analysis of fitted parameters given by the first two steps.

Steps one and two have been done in an automatic fashion *via* an *ad-hoc* MATLAB code named “u-Tubes”. (see Algorithm below). The first part of the code is devoted to the treatment of the data, including: smoothing of the signal, baseline correction, characteristics points detection, optical artifact detection and correction. For each curve, several observables are measured and stored, such as the slopes around zero force at contact and release, the slope of the tube (if any), the relaxation during the contact phase, and many others (≃140 in total). These measures were exploited for the direct estimation of the physical parameters and some of them served as guessing parameters for the fitting procedure. The most important task of this first part is the classification of each curve in the three main categories - “contact”, “detachement”, “tube” (either finite or “infinite”) - along with, for the tubes, the classification of the two type of discontinuities “rupture” and “slippage”. The second part of the code, settles the fitting of the data by means of the proper model related to detachment, “rupture” tube or “slippage” tube. The outcome of this part are the fitted parameters (*t*_1_, *t*_2_, *k*_1_, *k*_2_, *η*, *k*_1*N*_, *η_N_*, see Table 1) and a supplementary classification of the curves on their fitting “quality” based on the examination of fit convergence and magnitude of residuals.

Finally, the parameter values obtained in the two first steps were prepared for statistical analysis and representation using a set of Python scripts (with the use of Python dabest supplementary package, https://acclab.github.io/DABEST-python-docs/index.html, (4)). The details of each of these parts are presented below.

#### *μ*-Tubes algorithm

1. Read data (txt file from JPK)
2. Smooth data (moving average)
3. **procedure** Classification
4. Detect characteristic points (contact, wait, retraction)
5. Detect & Correct optical artefact
6. Measure geometrical parameters (forces, slopes, etc.)
7. Classify discontinuity: rupture, slippage
8. Classify curve: contact, adhesion, finite/infinite tube
9. Estimate mechanical parameters (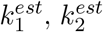, etc.)
10. **end procedure**
11. **procedure** Fitting
12. Assign model and constraints (depending on 8. & 9.)
13. Determine guessing values (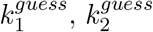 etc.)
14. Fit curve and get parameters (*k*_1_, *k*_2_, etc.)
15. Evaluate fitting quality from residuals
16. **end procedure**
17. Save and Plot data

### Curves processing

Raw curves were first smoothed with the built-in MATLAB function smooth (with a moving average algorithm). The next step of the data treatment was the spotting of characteristic time-force points coordinates.

### Characteristic points

These points mark a discontinuity in the force curve and delineate the boundaries for the curve segmentation in the three consecutive parts: contact, wait, retraction. In Fig. S3 (corresponding the panel A of Fig.2), from left to right, *t*_*c*1_ and *t*_*c*2_ are the time points at which the contact starts and ends respectively (in red); *t*_*r*1_ and *t*_*r*2_ are the start/end retraction points (in black); 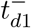 and 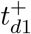 are the first discontinuity in the force curve during retraction (left and right limits), *t*_*d*2_ is the second discontinuity and, finally, *t*_*b*0_ is the time at which the force is back to zero amplitude. Notice that, 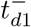 and 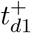 are coincident for a slippage rupture, *t*_*d*2_ is only defined for tubes, and 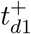 coincides with *t*_*b*0_ for adhesions. All these points, excepted those lying in the baseline, are detected finding the extremes of the second derivative of the time-force curve, using the MATLAB function `findpeaks` (Fig. S3A).

### Optical artefact correction

A typical force-curve should have a zero amplitude until the contact (*t* = *t*_*c*1_). However, some curves come with a positive amplitude for *t* < *t*_*c*1_ (~ 18% of the dataset), which has been understood as the signature of an optical artefact when part of the laser goes through the small T cell (see Fig. S3B, noised blue line between the two pink points). The detection of the artefact is based on a tolerance criterion of the max force-amplitude of this curve’s segment, fixed to the mean noise force amplitude 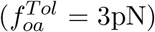. The correction is based on the assumption of the symetry around *t* = 0 when the push and pull velocities are equal, the contact force is moderate and the contact time is small. Once detected, the artefacts were smoothed, mirrored, shifted at *t*=0 (blue smoothed line), and subtracted to the original force-curve (cyan), which gives the final corrected signal (orange).

### Geometrical parameters

After the optical artefact correction, if any (see above), several geometrical parameters are then measured on each part of the curve (see Fig. S3A) and include contact/retraction slopes, force relaxation (*t*_*c*2_ ≤ *t* ≤ *t*_*r*1_), slope of the linear part (around *t* = 0), slope of the tube 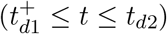, force-drop between 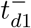 and 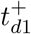, etc.. While the *t* < 0 part of the curve (in cyan, in Fig. S3) served for obtaining preliminary measures used as fitting guesses, the *t* > 0 part (in orange,) was exploited for fitting the model in Eq.2.

### Discontinuity classification

Two types of discontinuities are identified: “rupture” and “slippage”.

A rupture discontinuity is defined by the boolean defined by two logical conditions, on the absolute and relative value of the force drop at *t* = *t*_*d*1_ (if any):

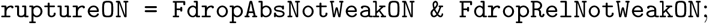

with the symbol & is the logical AND, and where

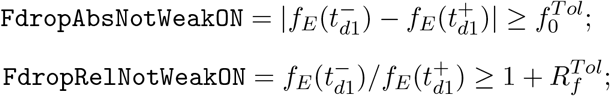

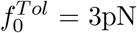 corresponds to the average peak-to-peak amplitude of the experimentally recorded noise on the optical tweezer data *f_E_*(*t*). We fixed 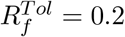.

Slippage discontinuities are simply defined by the boolean

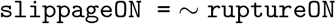

where ~ is the logical negation.

### Curve classification

Based on geometrical parameters measured on curves and on the characteristic points, four main categories of force curves are established: contact, detachment, finite tube, infinite tube. This classification is performed with the requirement of several logical conditions, which are all referred to the positive-time domain of the force curve. First, a curve is classified as a contact (ie. not showing any significant event upon separating the cell and the bead) if the maximum or the mean force over *t* > 0 are found smaller with respect to a multiple of the force tolerance 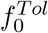. The logical condition is then

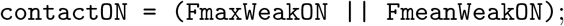

where the symbol || is the logical OR, and the two booleans are defined by:

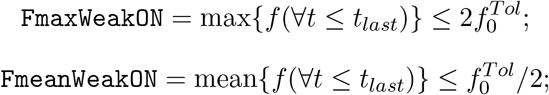

with *t_last_* – {*t*_*b*0_, *t_end_*}.

Second, a curve is classified as a detachment if the force amplitude drops to zero after the first event 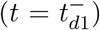, where the tolerance is now fixed on time, and corresponds to the minimum lifetime tolerance for tubes fixed to 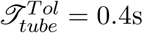 (equivalent to a maximum length of 1μm). The logical condition, with the respective tolerances, is

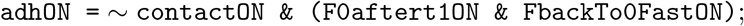

where

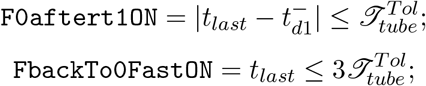

This implicatess that, even if a curve shows a tube-like fingerprint, it will be classified as a detachment if its lifetime is too short. The reason behind this choice is a matter of robustness: very short tubes are not very informative for performing a robust extrapolation of the tube parameters (*k*_1*N*_, *η_N_*), while they contains the information of the first elastic-like part of the model.

Finally, if none of the two previous conditions are trues, the curve is classified as a tube. The logical condition is then:

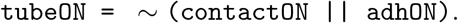

Tubes are subsequently classified as “finite” or “infinite”, where “infinite” tubes are essentially those lasting until the end of the experiment (until *t* = *t_end_*). In order to distinguish a tube from a residual weak force amplitude 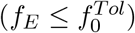, the mean force of the tube and his final force are verified to be bigger than the previous zero-force tolerance. The logical definition is the following

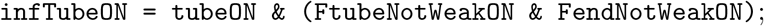

where

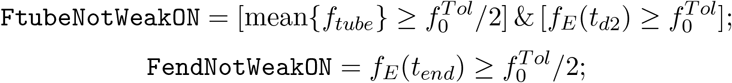

and

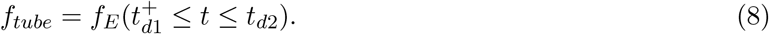

Finally, finite tubes are simply defined as

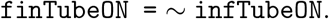

### Curve fitting

#### Parameter estimation

Several geometrical parameters were used for determining preliminary estimations of the mechanical parameters. First, we measured directly from the experimental force-curve *f_E_*(*t*), the slope at contact 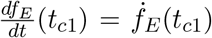 and at (negative force) retraction *ḟ_E_*(*t*_*r*1_), see Fig. S3, Inset, straight red and black lines, respectively. From these two slopes, we can obtain two estimations of *k*_2_ (which are similar, with typical differences due to small hysteresis in retraction (5)). For this, we assumed that the deformation of the molecule is negligible for both contact and negative retraction situations (*k*_1_ ~ 0, ∀*t* < 0). This is due to the fact that, differently from the pulling situation, pushing a single molecule from its equilibrium position leads to a negligible entropic contribution due to the molecular stiffness with respect to the pulling case.

Setting this condition in the model, and measuring the experimental force-slope of the negative retraction *ḟ_E_*(*t*_*r*1_), we get the following estimation of the cellular stiffness *k*_2_

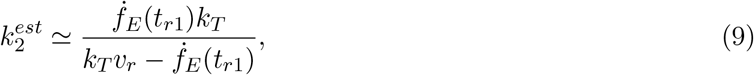

where we recall that *v_r_* is the retraction velocity of the piezo-electric stage, and *k_T_* is the stiffness of the optical trap. Obviously, only positive estimations where considered.

The same rationale conducted to the estimation of the whole elastic contribution of the system 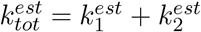, from the experimental force-slope of the positive retraction *ḟ_E_*(*t*_*r*2_) (Fig. S3, inset, straight blue line). In fact, for the positive retraction case, we assumed that also the molecule is loaded together with the membrane, contributing to the total elastic stiffness, which is then estimated as

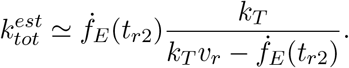

As a consequence, one can obtain an estimation of the receptor/cytoskeleton bond stiffness from

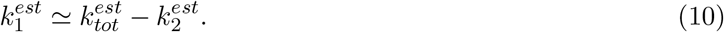

Moreover, the estimation of *k*_2_ allows to estimate *η* in the waiting segment (*t*_*c*2_ ≤ *t* ≤ *t*_*r*1_), by means of the direct fitting of the model with *k*_1_ ~ 0 since the system is not under traction (Fig.S3, Inset, orange curve). An approximation of *η* is then given by

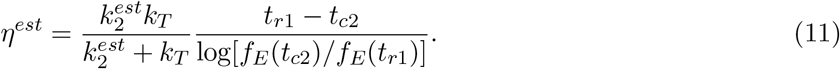

Important enough, as the total effective contact time *t_C_* = *t*_*r*1_ – *t*_*c*2_ is not the same for all the curves, we chose to estimate *η^est^* at *t_C_* = 0.4 sec for the entire the dataset.

From Eq.4, with the limit *t* → +∞, we can find the approximation of the time-force curve for long tubes 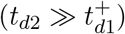. This gives the approximation of the experimental tube slope at his end, *ḟ_E_*(*t*_*d*2_), from which we get the estimation of the tube stiffness as

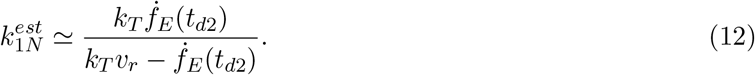

Finally, for long tubes the relaxation term of Eq.7, 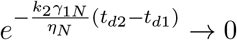, which lead to the approximation of the force value at end of the tube, ie. at *t* = *t*_*d*2_

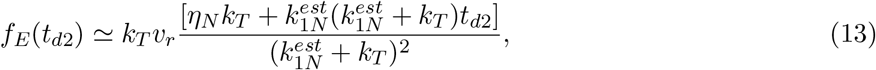

Considering 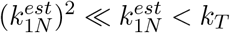, we get

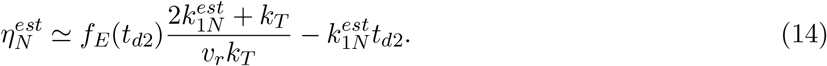

#### Guessing values

Overall, the guess values for the mechanical parameters *p^guess^* are fixed according to the estimated parameters *p^est^* if any, or to a prefixed value otherwise. In the latter case, the prefixed values have been arbitrary fixed to the median of the estimated parameters 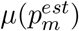, calculated over all the m curves for which *p^est^* exists. This case concerns only *η*_1_ and *k*_1*N*_, for which we have 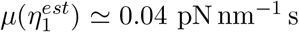, and 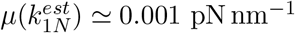. Accordingly, the guessing values are generally fixed to

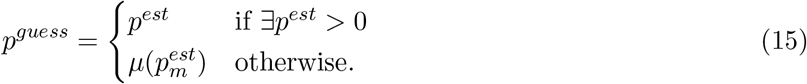

Note that this rule is slightly modified for curves exhibiting tubes for the two time-dependent parameters *k*_1_ and *η*, for which the rule becomes

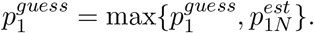

This condition guarantees that 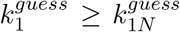 and 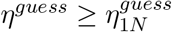, coherently with the fact that both stiffness and viscosity should not increase after the emergence of a tube These choices for the guessing values, even if not mandatory, increase the likelihood of a successful fit and consequently reduce the computational time.

Last, the guess value for the discontinuity event time is fixed to 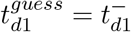.

#### Curve fitting

The fitting was performed by means of the MATLAB function `fmincon`, which find the minimum of a constrained nonlinear function. This function was used for minimizing the residual sum of squares (RSS) between theoretical and measured forces. Accordingly, the objective function has been defined as

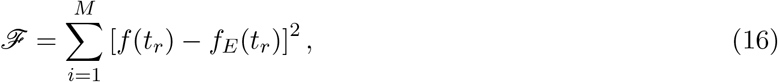

where 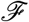 is homogeneous to a force, and *M* is the total number of points constituting the fitted force curve.

The minimization procedure has been subjected to various constraints, defined as linear or nonlinear combination of the free parameters *t*_*d*1_, *k*_1_, *k*_2_, *η*, *k*_1*N*_, *η_N_*. In particular, the constraints have been imposed on (i) both the slopes of the time-force curve at the origin and at the end of the tube, and (ii) the force amplitudes at time 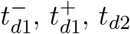 as follows.

First, the slope of the force at the origin of times, defined as 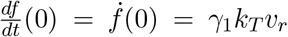, has been constrained to not differ by more than 10% from the experimental value of the slope at positive retraction *ḟ_E_*(*t*_*r*2_), hence:

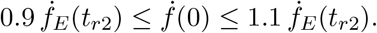

The constraint on the slope of the tube was

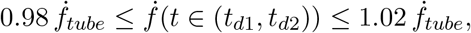

where *f_tube_* is defined in Eq.8. Second, the constraints on the forces at the first discontinuity have been fixed to

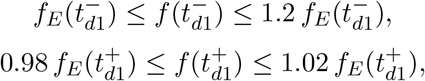

and at the second discontinuity was

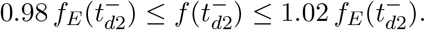

For the maximum force peak before transition, 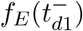, we fixed a bigger tolerance with respect to the other values because a small subset of curves present a fast change on the force-slope preceding the discontinuity at 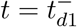. This change does not correspond to the relaxation term introduced by the viscous dashpot, and - for preserving simplicity - we choose to not account for this (occasional) behaviour.

Finally, the discontinuity in 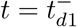 has been modeled as a “degradation” of both the molecular elastic (*k*_1_) and cellular viscous (*η*) parameters, for which we imposed that *k*_1*N*_ ≤ *k*_1_ and *η_N_* ≤ *η*.

To avoid potential non-physical solutions, we defined a set of lower and upper bounds (*lb*, *ub*) for all the parameters, so that the fitting solution of a parameter *p* is always in the range *p^lb^* ≤ *p* ≤ *p^ub^*. For the majority of the parameters, we fixed the lower/upper bounds to very small/big values (see Table 2) with respect to their final median (see Table 1, main text).

**Table 2:**
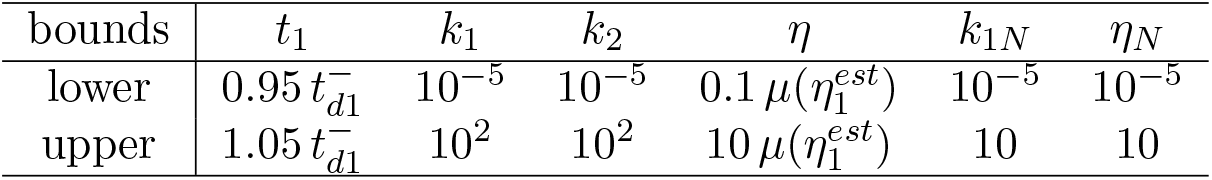
Upper and lower bounds for each fitting parameter

For the particular choice of *t*_1_ and *η* bounds, we did as follow. First, we limited *t*_*d*1_ to a very narrow region around the point spotted on the curve 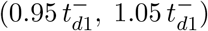, the transition being generally well identified for the majority of force curves. Second, we limited *η* to the region around the median of its estimated value 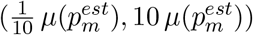, due to the large variance of the corresponding distribution. This has two counterparts: from the one hand, it makes risky to fix *η* exactely to its median; on the other hand, too small or too big values of *η* can lead to a failure of the fitting algorithm. The great variance of *η^est^* reflects the difficult to extrapolate this parameter, which is mostly related to the quasi-linear behaviour of the majority of the force-curves where the term 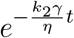 approaches zero.

#### Fit quality and residuals

The difference between the experimental and the fitted curve has been evaluated in term of the residual standard error (RSE), obtained by taking the square root of the objective function of Eq.16 normalized by *M* – 2. The RSE was separately evaluated before and after the first discontinuity (corresponding to 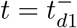) such as

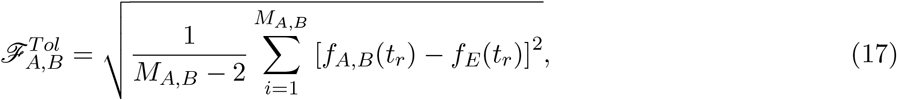

where, *f_A_*(*t*) and *f_B_*(*t*) correspond to Eq.6 and Eq.7, respectively. Accordingly, *M_A_* and *M_B_* represent the number of points of the force curve for 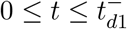 and 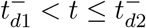.

Based on the RSE values, we evaluated the fit quality of each curve with the following boolean

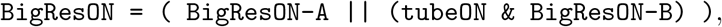

where

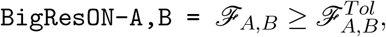

with 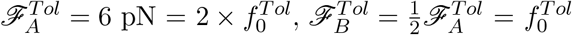.

### Data post processing

The post processing of the data is based on the following python librairies, mainly avalaible in Anaconda Python Distribution (https://anaconda.org/; numpy, scipy, scikit, pandas) or on the web (dabest; ttps://acclab.github.io/DABEST-python-docs/index.html (4)). The visualisation has been made with matplotlib, seaborn (https://seaborn.pydata.org/index.html) and dabest packages.

#### Data preparation, sorting and cleaning

We first loaded the data output by the fitting and classification procedure as .xls, from the different experimental sets, and curated it for easy further processing. We then used booleans present in the data file to remove curves having been labelled by the entire procedure as rejected (eg. because of too large fitting residuals).

Comparing at that stage the differents data sets (slightly different *k_T_*, contact forces 10-15pN) we observed that in the experimental ranges, neither the dispersion of the fitted parameters nor the central tendancies depend on the initial data setdata. We then confidently pooled all data sets in further analysis.

We then subsetted the data to short contact times, between 0 and 0.5 sec, to be sure to have mainly unique tubes in our analysis.

From the fitting strategy we presented in the relevant section, the following pooling of data have been made [see Table 1 in the main text]. The fit has been faithfully estimating *k*_2_ for rupture and slippage tubes; *k*_1_ for rupture tubes only, *k*_1*N*_ for rupture and slippage tubes; eta for rupture and slippage tubes; etaN for rupture and slippage tubes. This allowed us to plot, separating each antibody case with or without latrunculin, the final population of acceptable values. We present the obtained data sets in the Fig. 3 in the main text and in the SI, in particular Fig. S5.

#### Data representation and statistical tests

We chose to use a Data Analysis with Bootstrap-coupled ESTimation strategy (dabest Python package) (4).

We set to evaluate (a) the relative variations of the parameters estimated without latrunculine among the different antibodies used as handles to pull adhesion events or tubes to detect the molecule effect on the different mechanical parameters, and (b) the relative variation, for each parameter and antibody, of the value with vs. without the drug presence as an indicator of the cytoskeleton on each parameter, for each molecule.

This methodology allows to represent the dataset explicitely and uses a bootstraping approach to estimate the distribution of the differences between two sets of data (eg. between without and with latrunculin for a given parameter and a given molecules) or between one reference and other data sets (eg. Comparing aCD3 to each of the others antibodies).

The estimation plot produced allows to conclude if, for a given CI value (here 95%), data sets are extracts of different or not populations. Where a data set was observed to be significantly different (in terms of dabest analysis) from its comparison / reference distribution, we indicated it on the graphs by a star symbol (*).

#### Pooling detachment and tube data

Mechanical parameters can be obtained in principle from the detachment curves ()see Table 1 in the main text), but with a reduced accuracy, in particular for *k*_1_. This is illustrated on Fig. S6, where the dispersion can be appreciated. This dispersion implicates that some of the significant differences observed for the tubes only data are affected, but not the relative variations of their median values. As a consequence, we did not pool the detachment data with the Rupture case.

## Supplementary Figures

**Figure S1:**
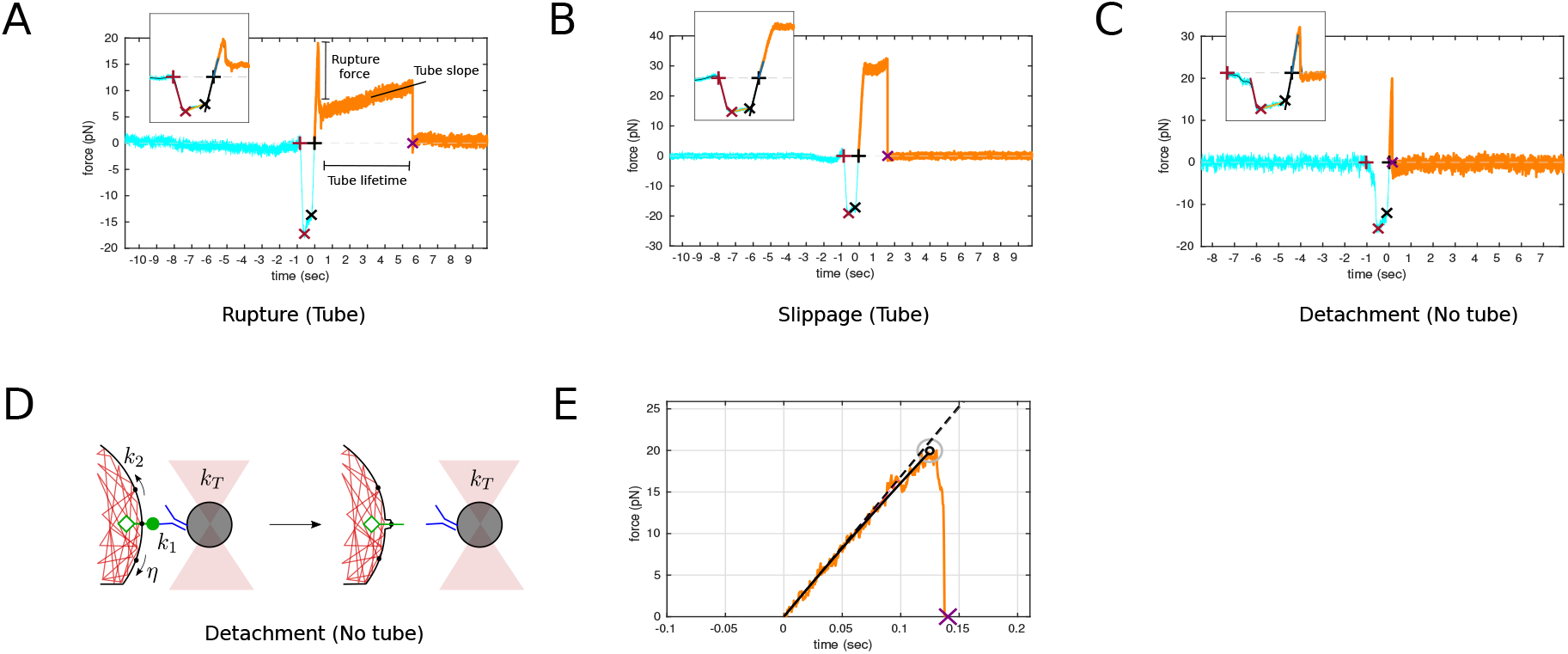
Typical force vs. time curves: A. Rupture case, tube. B. Slippage case, tube. C. Detachment case (no tube). Insets are presenting zooms over the contact region. D. Model schematics for detachment. E. Fitting model to the detachment data presented in C, as done for the rupture and slippage cases in main text Fig. A,B. Note that no typical”contact” cases is shown here since it is not bringing relevant information to the present study.

**Figure S2:**
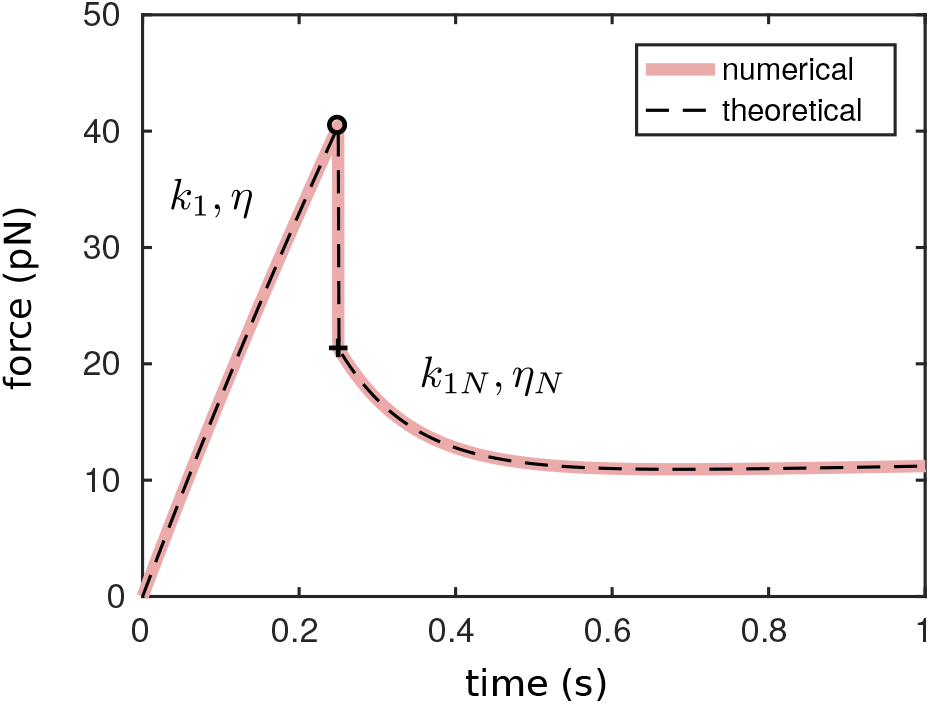
Theoretical results of the measured force *f*(*t*) of a membrane tube extrusion experiment, both numerical (red) and analytical (dotted-black), with *x*_0_ = 0nm, *v_r_* = 2.5μms^-1^, *t_r_* = 0 sec, and *f_0_* = 0 pN. The force history is caracterized by two regimes: for 0 ≤ *t* ≤ *t_d_* an almost linear regime followed by a very moderate relaxation, at *t* = *t_d_* an instantaneous release of the force, due to the abrupt change in stiffness *k*_1_ → *k*_1*N*_, and - finally - for *t* > *t_d_* a second relaxation followed by a quasi-plateau of the curve.

**Figure S3:**
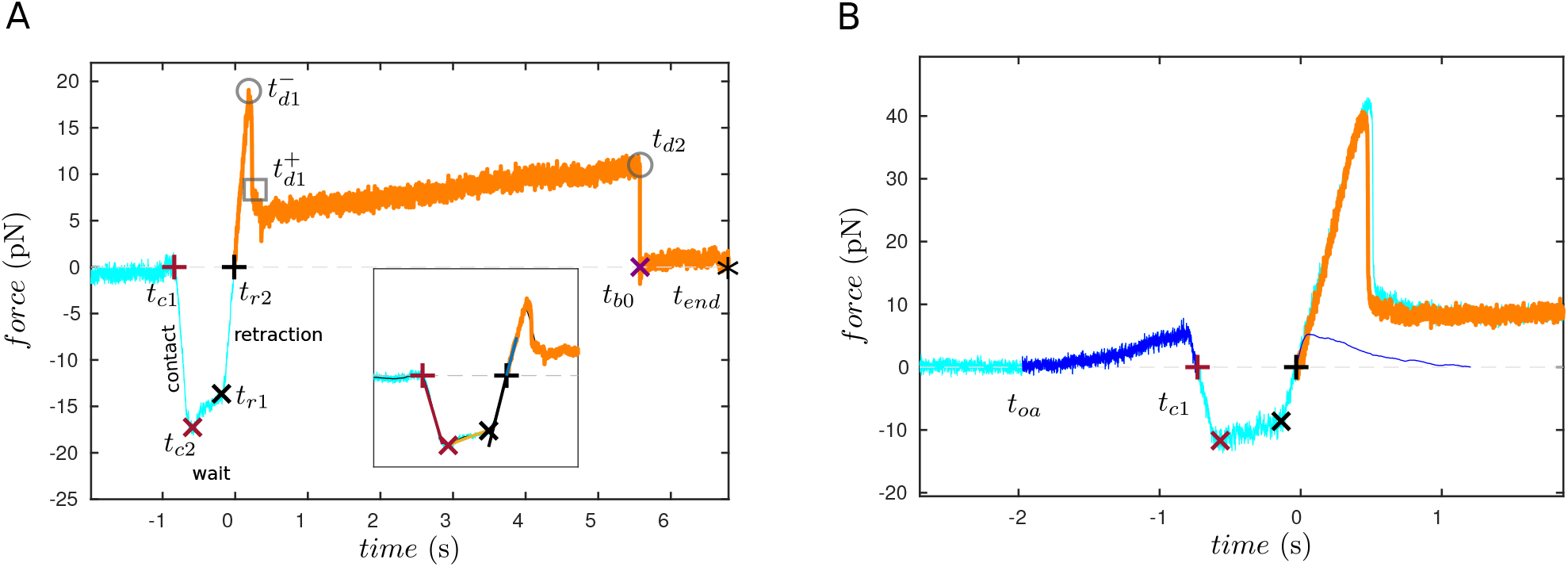
Experimental force-curve for a “rupture” event. (A) Characteristic points spotted on a typical curve. *t*_*c*1_ and *t*_*c*2_ are the time points at which the contact starts and ends respectively (in red); *t*_*r*1_ and *t*_*r*2_ are the start/end retraction points (in black); 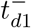 and 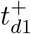 are the first discontinuity in the force curve during retraction (left and right limits), *t*_*d*2_ is the second discontinuity and, finally, *t*_*b*0_ is the time at which the force is back to zero amplitude. Notice that, 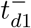 and 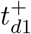 are coincident for a slippage rupture, *t*_*d*2_ is only defined for tubes, and 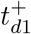 coincides with *t*_*b*0_ for detachment curves. (B) Optical effect correction. Light blue, original data; dark blue data between *t_oa_* and *t*_*c*1_ optical effect on the pressing segment of the curve, average and symetrized for the pullingon segment, blue thin line; orange, corrected data on pulling segment.

**Figure S4:**
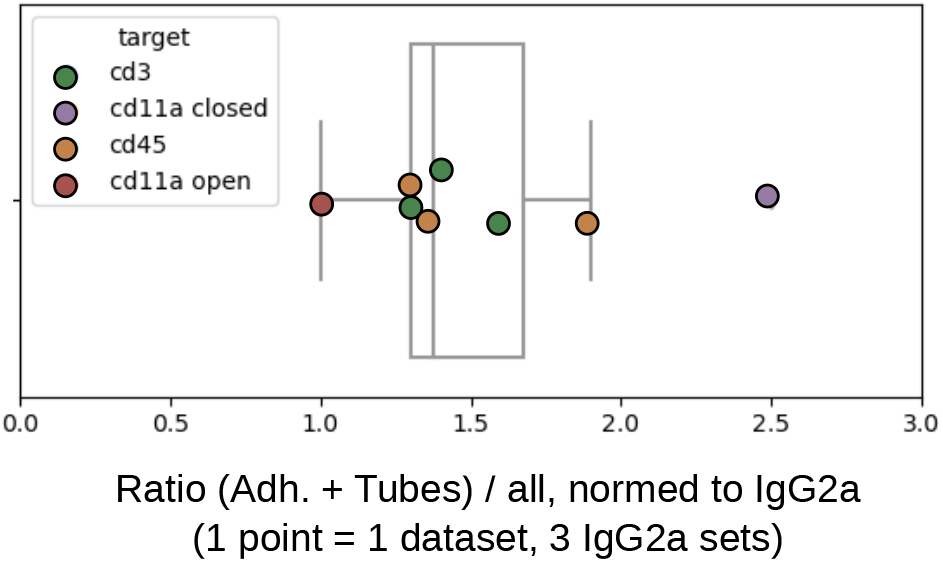
Specificity of the different antibody handles, compared to IgG2a isotype control. The graph presents the ratio of (adhesion+tubes) to the total number of curves, per handle molecule. Note that since our Jurkat T cells are not activated, the number of interactions that was recorded with the antibody directed toward the open conformation of LFA1 was low, and lower than for the closed state of this integrin (6).

**Figure S5:**
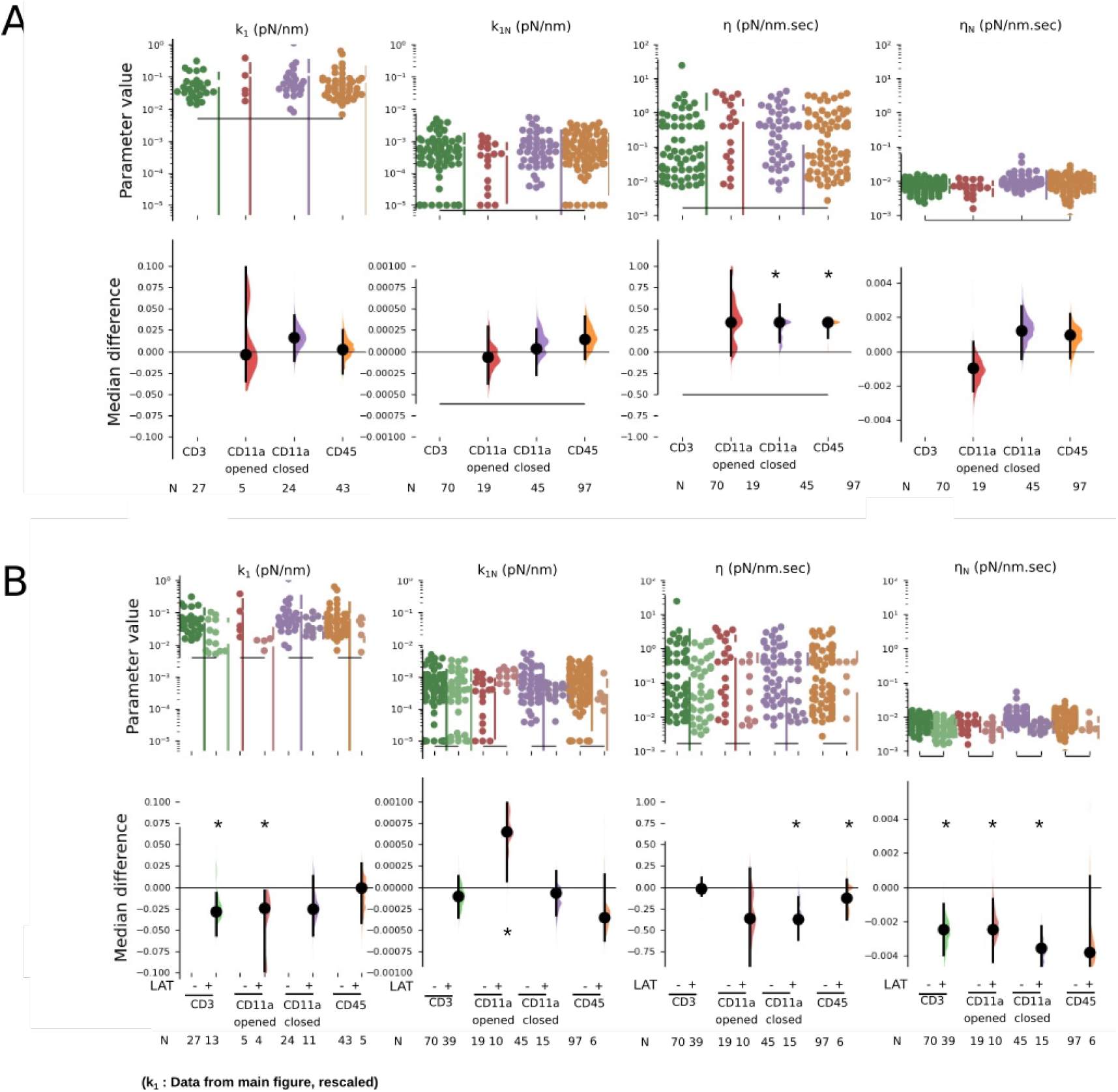
A. Estimation plots for *k*_1_ (reproduced from main text data), *k*_1*N*_, *η* and *η_N_*, for all antibody handles, and relatively to CD3 as a reference, without LatA treatment. B. Estimation plots for the same parameters, including the data where latrunculine was added. Here, the comparison is made between the cases without and with the drug, for each handle. One point corresponds to one fitted curve.

**Figure S6:**
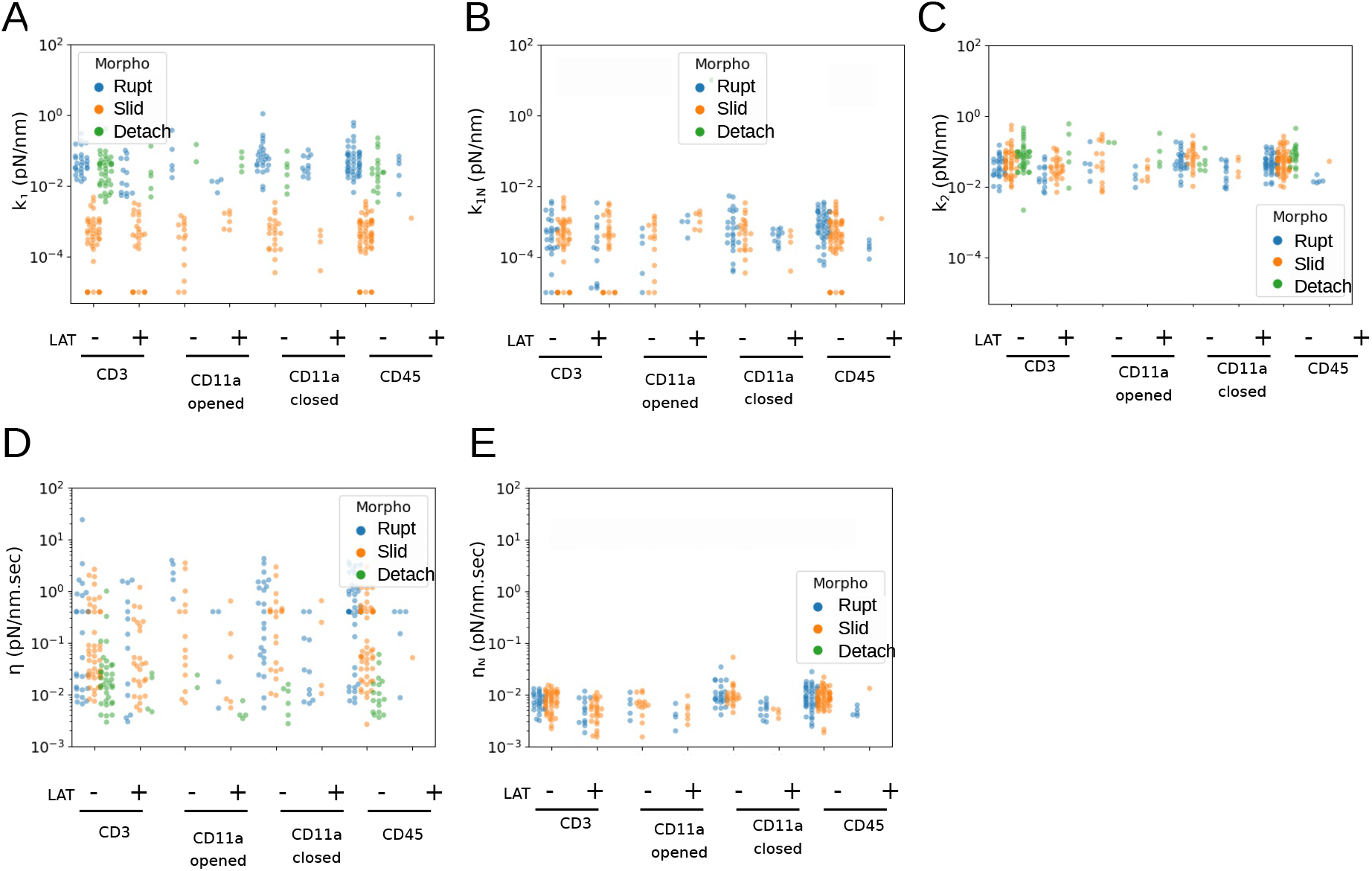
Scatter plots of all mechanical parameters, extracted from the experimental data, as a function of the antibody handle, presence or not of LatA treatment and morphology (Rupt = “rupture” tube, Slip = ‘‘slippage” tube, Detach = detachment). Please note that the *k*_1*N*_ and *η_N_* values for adhesion curves are not existing by model definition. One point corresponds to one fitted curve.

**Figure S7:**
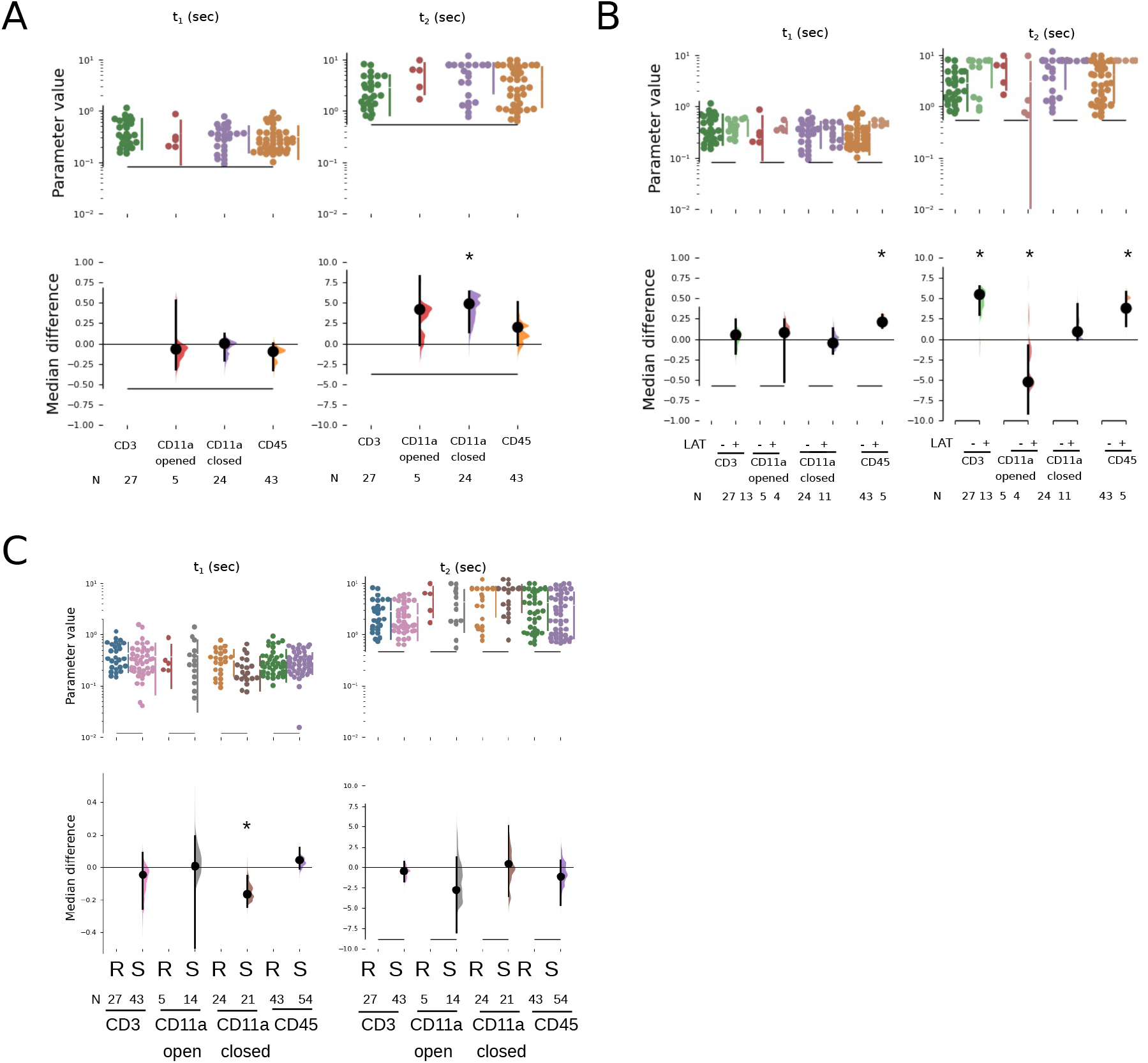
A. Estimation plots for *t*_1_ and *t*_2_, taking CD3 as a reference, for “rupture” tubes only. B. Estimation plots for the same parameters, including the data where latrunculine was used. Here, the comparison is made between the cases without and with the drug, for each antibody handle. C. Comparison, per antibody handle, between “rupture” (R) and ‘‘slippage” (S) tubes. One point corresponds to one fitted curve.

**Figure S8:**
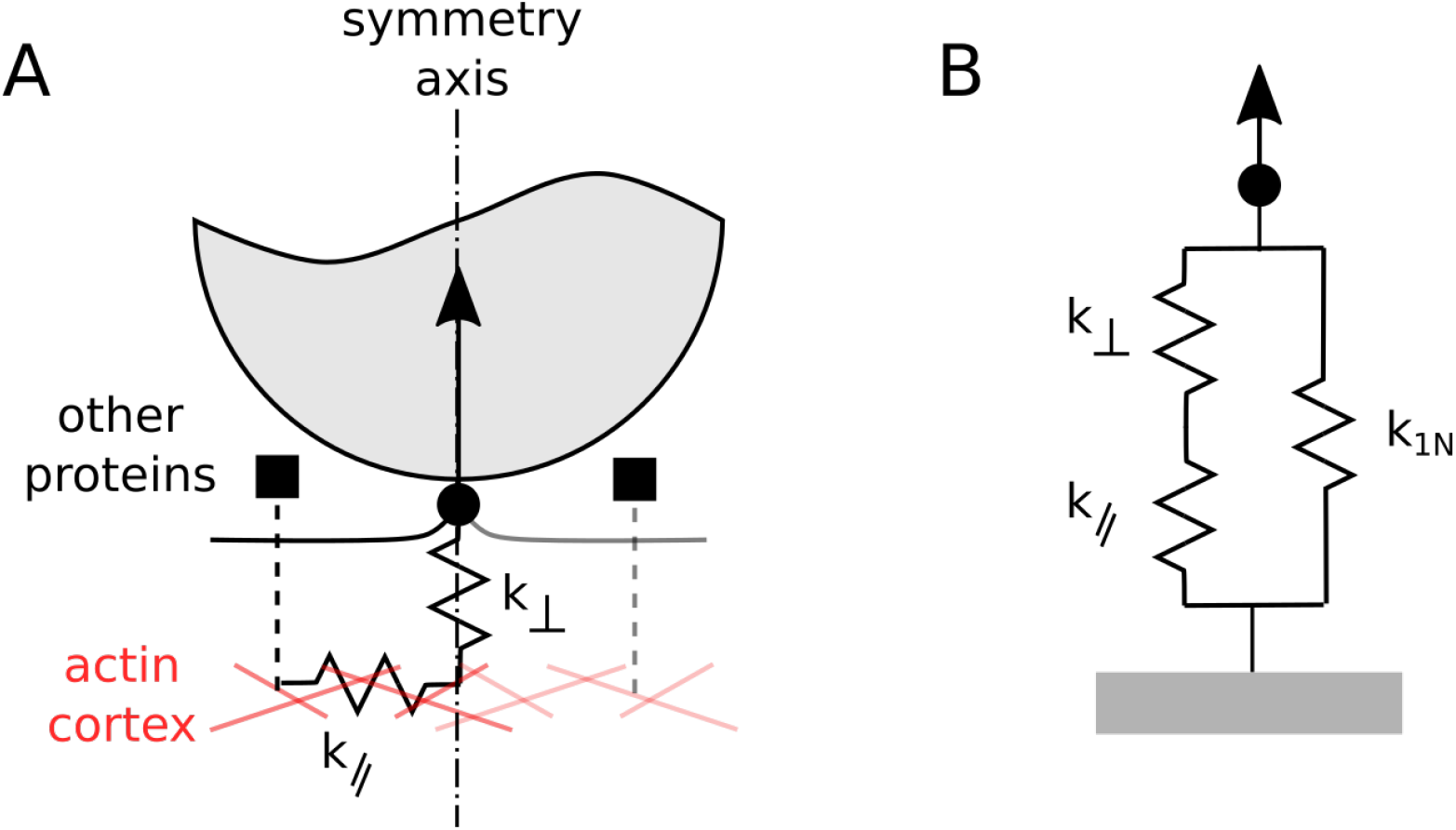
Schematics of the early moments of tube pulling. B: Equivalent spring-based model

**Figure S9:**
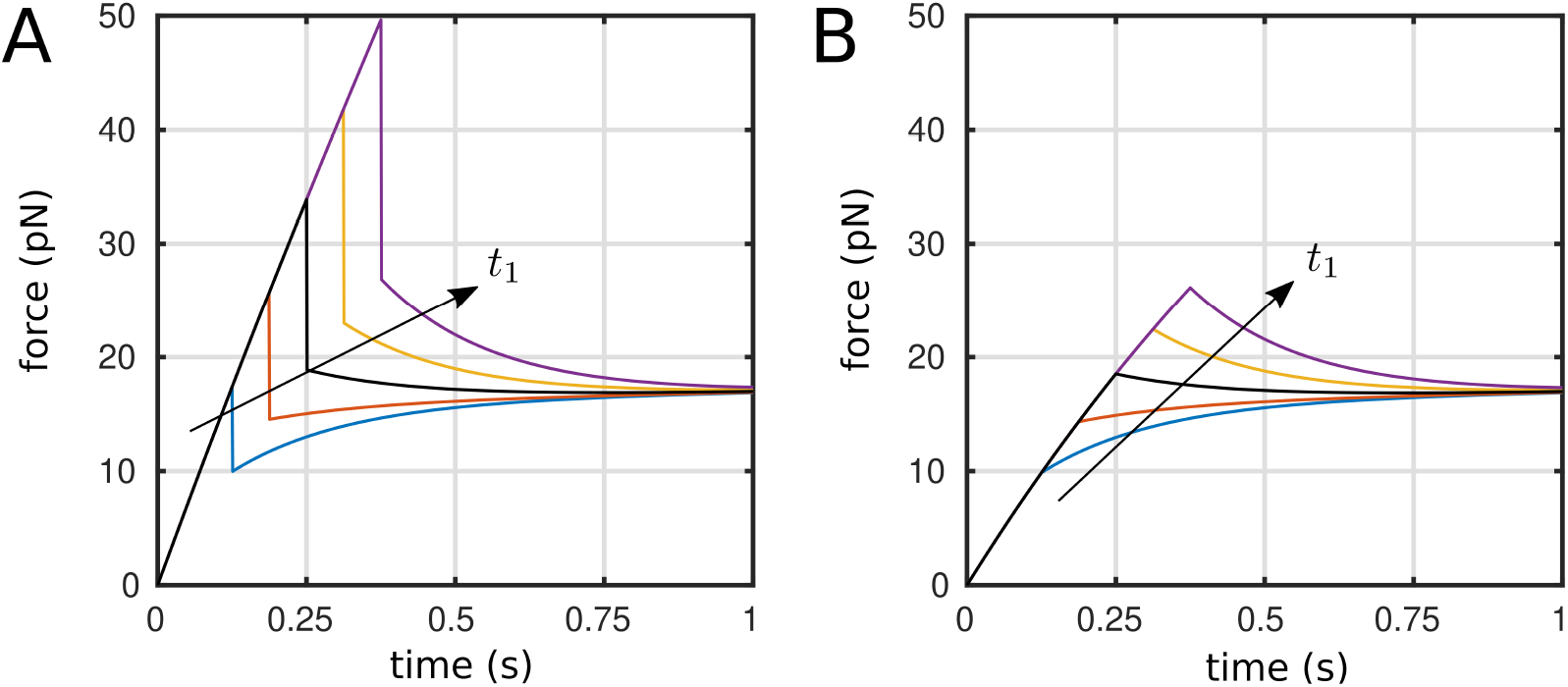
Parametric study of the viscoelastic model for the time of the discontinuity *t*_1_. A: Rupture case. B: Slippage case. The black curve in between the others correspond to the one obtained via the fitting in Fig. S1, left panel. Parameters are *v_r_* = 2000nms^-1^, *k_T_* = 0.25pN nm^-1^, *k*_1_ = 0.05pNnm^-1^, *k*_2_ = 0.05pNnm^-1^, *η* = 0.04pN nm^-1^ s, *k*_1*N*_ = 0.0005pNnm^-1^, *η_N_* = 0.008pNnm^-1^ s, and *t*_1_ = 0.25s. The others curves have been obtained multipling the value of t1 by the following vector of factors {0.1, 0.5, 1, 2, 5}.

**Figure S10:**
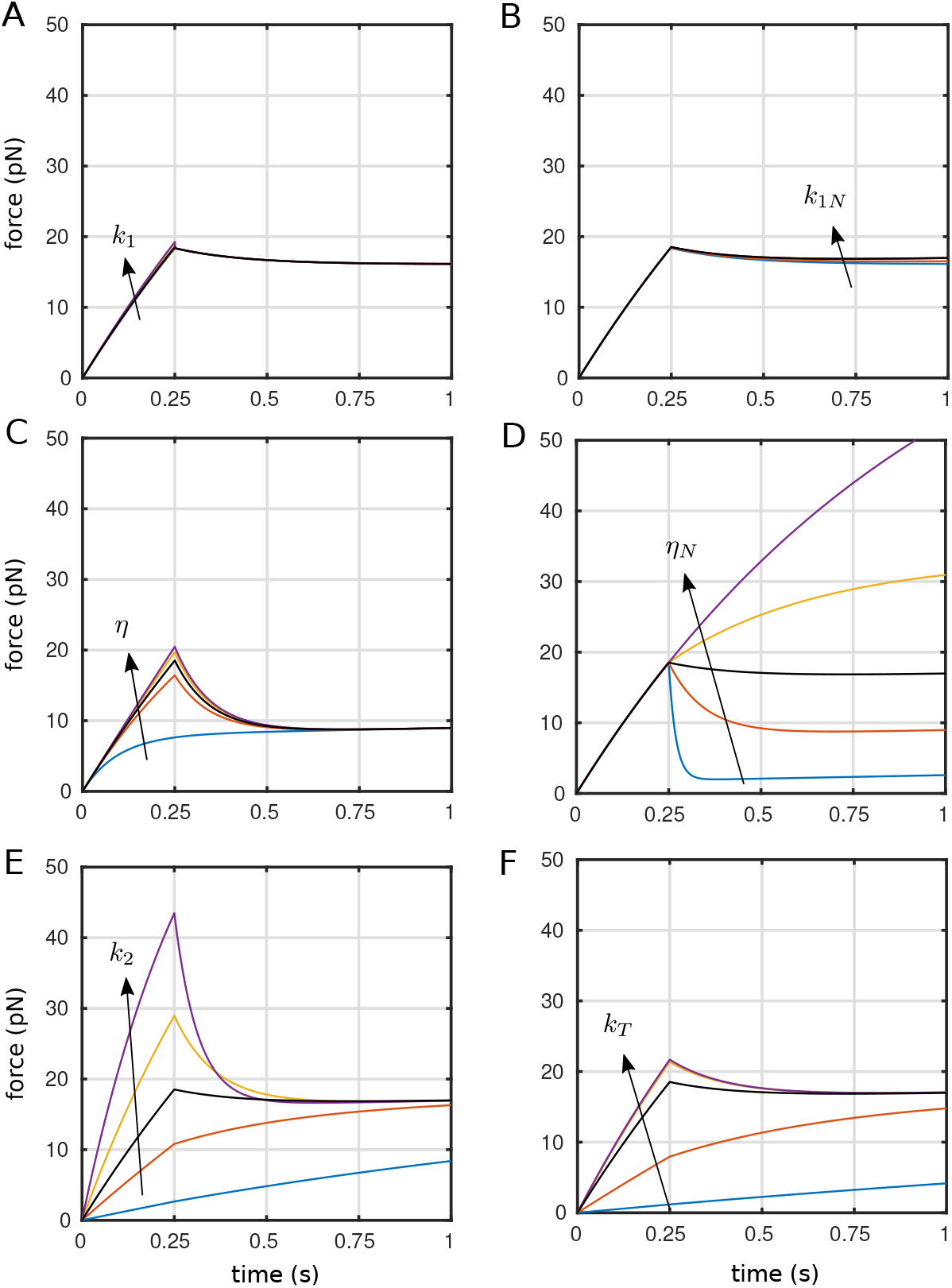
Parametric study of the viscoelastic model for the slippage case. The black curve inbetween the others correspond to the one obtained via the fitting in Fig. S1, left panel. For panels from A to F, parameters are *v_r_* = 2000nms^-1^, *k_T_* = 0.25pN nm^-1^, *k*_1_ = *k*_1*N*_ = 0.0005pNnm^-1^, *η_N_* = 0.008pNnm^-1^ s, *k*_2_ = 0.05pN nm^-1^, *η* = 0.04pN nm^-1^ s, and *t*_1_ = 0.25s. The others curves have been obtained multipling these values by the following vector of factors {0.1, 0.5, 1, 2, 5}. For panel F, parameters are *k_T_* = 0.01, 0.1, 1, 10, 100

